# Cryo-electron tomography reveals how COPII assembles on cargo-containing membranes

**DOI:** 10.1101/2024.01.17.576008

**Authors:** Euan Pyle, Elizabeth A. Miller, Giulia Zanetti

## Abstract

Proteins traverse the eukaryotic secretory pathway through membrane trafficking between organelles. The COPII coat mediates the anterograde transport of newly synthesised proteins from the endoplasmic reticulum, engaging cargoes with a wide range of size and biophysical properties. The native architecture of the COPII coat and how cargo might influence COPII carrier morphology remain poorly understood. Here, we have reconstituted COPII-coated membrane carriers using purified *S. cerevisiae* proteins and cell-derived microsomes as a native membrane source. Using cryo-electron tomography with subtomogram averaging, we demonstrate that the COPII coat binds cargo and forms largely spherical vesicles from native membranes. We reveal the architecture of the inner and outer coat layers and shed light on how spherical carriers are formed. Our results provide insights into the architecture and regulation of the COPII coat and advance our current understanding of how membrane curvature is generated.

## Introduction

Eukaryotic cells utilise the secretory pathway to transport proteins and lipids to their required locations within and outside the cell. Approximately one in three proteins are translocated in the endoplasmic reticulum (ER) upon synthesis and are trafficked to the Golgi apparatus as the first step of the secretory pathway^1^. Anterograde transport of proteins from the ER to the Golgi is facilitated by coat protein complex II (COPII)-coated membrane carriers. The COPII coat assembles on the cytosolic side of the ER membrane, generating membrane curvature to form coated carriers, while specifically recruiting and enveloping newly-synthesised cargo proteins^2,3^.

COPII comprises 5 proteins (Sar1, Sec23, Sec24, Sec13 and Sec31) that are essential and highly conserved from yeast to humans^3^. COPII assembly is initiated by the small GTPase Sar1, which inserts its N-terminal amphipathic helix into the outer leaflet of the ER upon nucleotide exchange, an event catalysed by the ER-resident GTP Exchange Factor (GEF) Sec12^4,5^. Membrane-bound Sar1 recruits heterodimeric Sec23/Sec24 to form the inner layer of the COPII coat, with Sec24 acting as the main cargo-binding subunit^6,7^. The outer coat layer is formed when heterotetrametric rod-shaped Sec13/Sec31 complexes are recruited to budding sites, via interaction of Sec31 with Sec23/Sar1, and assemble in a cage-like arrangement^8–10^. Symmetric polyhedral cages assemble *in vitro* when purified Sec13-Sec31 heterotetramers are incubated in the absence of any membrane^10,11^. The detachment of Sar1 from the membrane is triggered by GTP hydrolysis, stimulated by its cognate GTPase Activating Protein (GAP) Sec23, and further accelerated by binding of Sec31^12^. Sar1 GTP hydrolysis is thought to destabilise the coat; however, the dynamics and regulation of coat disassembly are poorly understood.

Previously, determined the structure of the *S. cerevisiae* COPII coat reconstituted *in vitro* from giant unilamellar vesicles (GUVs) using cryo-electron tomography (cryo-ET) with subtomogram averaging (STA)^13–16^. We showed that COPII forms coated tubes on GUVs and that the inner and outer coat layers both arrange into pseudo-helical lattices that wrap around the tubular membrane. High-resolution STA yielded atomic models describing coat interactions that allowed us to design coat mutants where assembly interfaces are disrupted^15^. We found that the two interfaces that form the outer coat cage, formed by the N- and C-terminal domains of Sec31, are dispensable for membrane budding *in vitro* and in yeast cells lacking the GPI-anchored protein cargo adaptor Emp24^15,17^. Moreover, when the interface between inner coat lattice subunits was weakened by mutation, budded membranes switch from a tubular to a spherical profile, indicating that membrane curvature is generated by a complex network of interactions spanning both coat layers^15^.

COPII-coated membrane carriers are known to adopt a range of sizes and shapes, which may be important to adapt to the wide range of cargoes that need to be accommodated. However, it remains unclear how coat assembly is regulated to achieve a variety of membrane carrier sizes^3,18,19^. Whilst our previous studies found that purified *S. cerevisiae* COPII forms extended tubules on GUVs, electron microscopy studies of cell sections suggest that membrane carriers *in vivo* are spherical vesicles 50-100 nm wide^20–22^, raising the question of which components of native membrane composition affect coat assembly and budding morphology. It also remains unclear how the tightly packed inner coat assembly is compatible with cargo binding by the Sec24 subunits. To answer these questions, we carried out *in vitro* reconstitution of COPII budding using native ER membrane derived from yeast, referred to as microsomes. In striking comparison to the tubules COPII forms on GUVs, cryo-ET revealed that the majority of coated membranes are pseudo-spherical. We used STA^16,23,24^ to obtain the structures of the inner and outer coat assembled on native membranes. We found that the inner coat layer can assemble as in its tubular arrangement, but forms limited patches of coat that are randomly oriented around a pseudo-spherical membrane. Cargo density could be detected within the inner coat array, in the space between inner coat subunits, indicating that lateral assembly of Sar1-Sec23-Sec24 heterotrimers can occur while small or flexible cytosolic domains of cargo molecules are accommodated in between. Finally, subtomogram analysis of the outer coat layer revealed non symmetric cages with a variety of architectures. We characterise multiple points of flexibility which were not previously described, increasing the complexity of the outer coat network, and challenging the current model where assembly of the outer coat into a polyhedral cage is the main driver of membrane curvature.

## Results

### COPII Forms Coated Pseudo-Spherical Vesicles on Microsomes

In order to reconstitute COPII budding *in vitro* from native membrane sources, we incubated purified *S. cerevisiae* COPII proteins with *S. cerevisiae* ER-enriched microsomes and a non-hydrolysable GTP analogue (GMP-PNP) (Figure 1A and Extended Data Fig. 1A). Imaging these budding reactions using cryo-ET revealed that COPII primarily forms vesicles (96.3 % of all coated membranes) on microsomal membranes which are clearly coated with both the inner and outer coat (Figure 1B). Only a minority of coated tubules were observed (3.7%), in striking comparison to previous reconstitutions using GUVs (91.4% tubules) (EMPIAR-11257). Most vesicles appeared fully coated (Figure 1A and Extended Data Fig. 1B).

**Figure 1.**
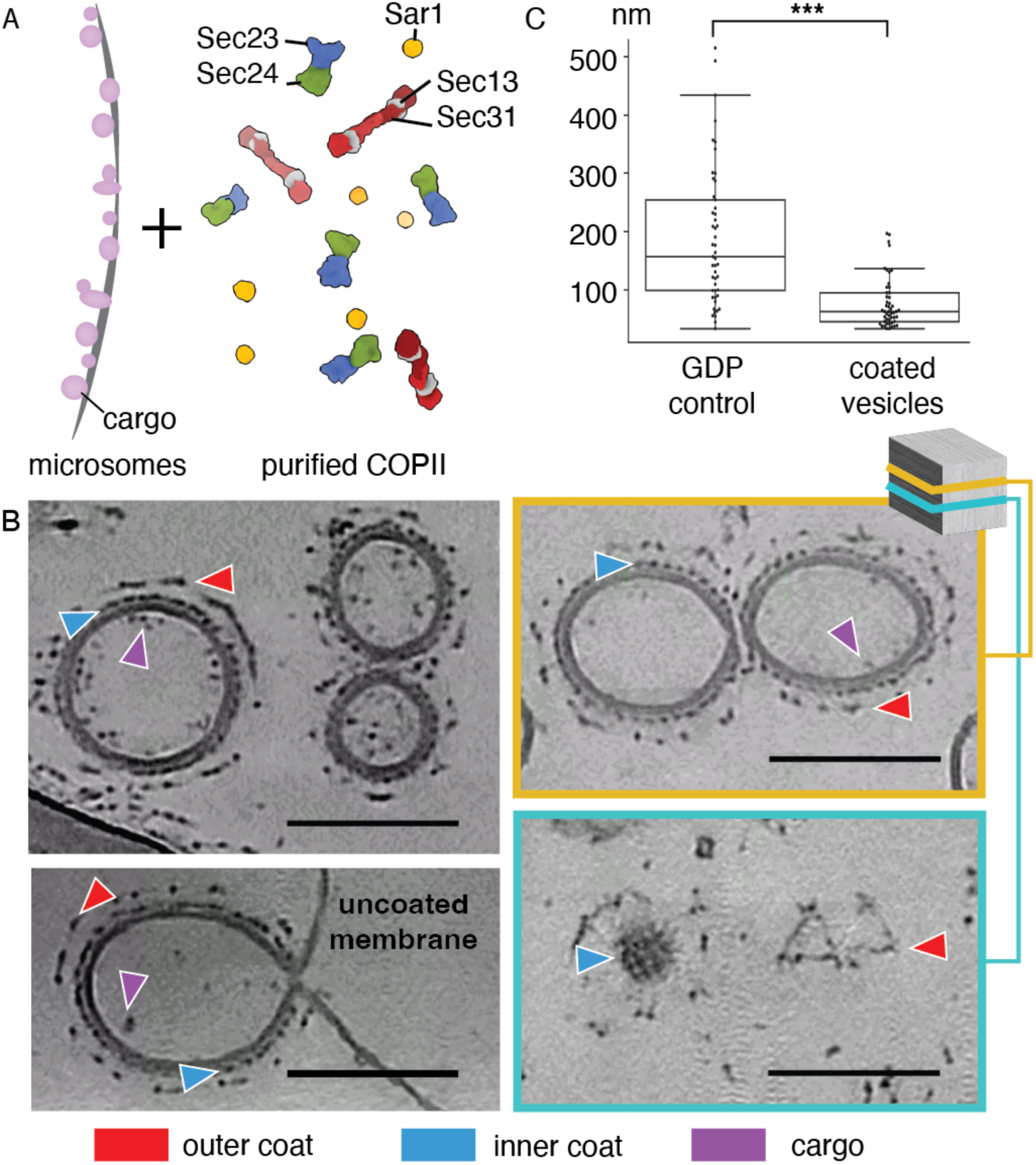
(A) Schematic of the in vitro reconstitution approach. (B) XY slices of representative reconstructed tomograms where instances of inner coat, outer coat and cargo are labelled with blue, red and purple arrowheads respectively. In the bottom left panel an example of a vesicle connected to its origin membrane via a neck is shown. The right panels represent two Z slices of the same tomogram. The bottom slice shows both the inner and outer coat layers of neighbouring vesicles. Scale bar = 100 nm (C) Membrane diameters were measured from a control reconstitution reaction where GDP was used and compared with the diameters of coated membranes obtained in the presence of GMP-PNP (n=1). Box plots centres, boundaries and whiskers represent median, 25 and 75 percentiles, and minimum and maximum values within box boundaries + 1.5X interquartile range, respectively. Statistical significance was determined using a homoscedastic two-tailed t-test (p=1.7E-08).

The microsome-derived COPII coated vesicles are significantly smaller than the donor membranes, as measured in a control sample where GDP was supplemented in place of GMP-PNP, demonstrating that the membrane is being actively deformed by COPII (Figure 1C). Most vesicles were detached, with only a handful of instances where coated vesicles were connected to other membranes via a constricted neck (Figure 1B and Extended Data Fig. 1C). Given that we used non-hydrolysable GTP analogues and performed no centrifugation or other mechanical perturbation of the sample, this suggests that vesicle scission from donor membranes may not depend on GTP hydrolysis.

### The Inner Coat Lattice Assembles in Small Patches on Vesicles

Previous high-resolution STA structures of GUV-derived tubules showed that the inner coat assembles laterally to form a pseudo-helical lattice^15,16^. To assess if and how the previously characterised assembly interfaces can give rise to spherical vesicles, we used STA to obtain a structure of the inner coat on vesicles (Figure 2A and 2B). We found that the arrangement of a subset of the inner coat is analogous to that previously described on tubes, with Sar1-Sec23-Sec24 trimers assembling laterally and longitudinally in an ordered lattice (Figure 2A-C, Extended Data Fig. 1D). At the resolution obtained (14.5 Å), there were no noticeable differences in the overall structure of the inner coat between the vesicles and the tubes, aside from the underlying membrane having spherical rather than tubular curvature. Consequently, we could unambiguously fit a previous high-resolution structure (PDB: 8BSH) of the inner coat into the density. However, the overall arrangement of the inner coat lattice differs significantly. On spherical vesicles, the inner coat lattice forms in small patches (Figure 2C). These patches can be orientated in different directions to one another on the same vesicle, suggesting that separate inner coat arrays can co-exist at multiple sites on the vesicle surface (Figure 2C). Visual inspection of tomograms shows that vesicles appear to have an inner coat even where patches are not detected, indicating that ordered patches and un-ordered individual subunits co-exist on spherical membranes (Extended Data Fig. 1D).

**Figure 2.**
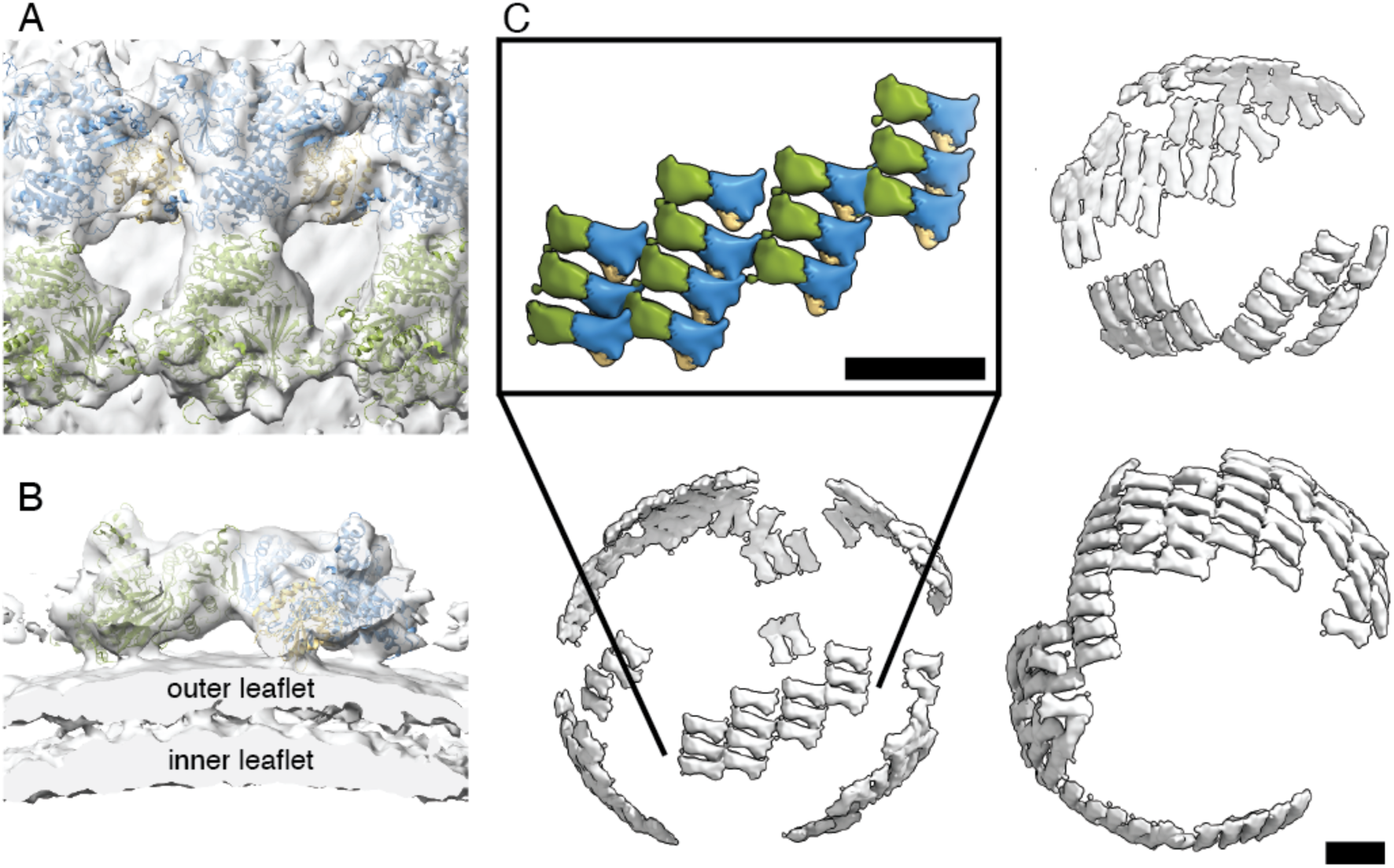
(A and B) Subtomogram average of inner coat on vesicles fitted with three copies of the Sec23-Sec24-Sar1 heterotrimer atomic model (PDB: 8BSH) with Sar1 in yellow, Sec23 in blue, and Sec24 in green. View in (A) is looking down towards the membrane (top view), while (B) cuts through the membrane (side view). (C) A low-pass filtered STA structure is mapped back in space. Three examples are shown to demonstrate the small patches arrangement of the lattice. Inset: a close up of one of the patches with the same colour code as in (A). Scale bar= 10 nm.

### Cargo Binds Within the COPII Inner Coat Lattice

We next set out to establish whether inner coat lattice formation is compatible with the presence of cargo. The inner coat is known to bind to a range of cargo molecules on several previously characterised binding sites on Sec24, including: the A-site located on the Sec24 side distal to Sar1 within the heterotrimer, and the B-, C- and D-sites located closely to one other on the opposite face of Sec24^25,26^. If cargo is bound to Sec24 in our structure, we would expect to see extra protein density proximal to the known binding sites. As we were unable to visualise density clearly above noise levels, we calculated the difference map between our STA structure of the inner coat on microsomes and a map generated by low-pass filtering the fitted model of the Sar1-Sec23-Sec24 heterotrimer to 14.5 Å. From the difference map, we found strong signal in the space between neighbouring Sec24 subunits, indicative of the presence of protein density and thus, potentially cargo (Figure 3A). The difference density seems located closest to the B-, and C-cargo binding sites of Sec24. As a control, we repeated the same experiment using the previously determined structure of the inner coat on cargo-less GUVs^16^, for which the difference map appears clear of any density (Figure 3B).

**Figure 3.**
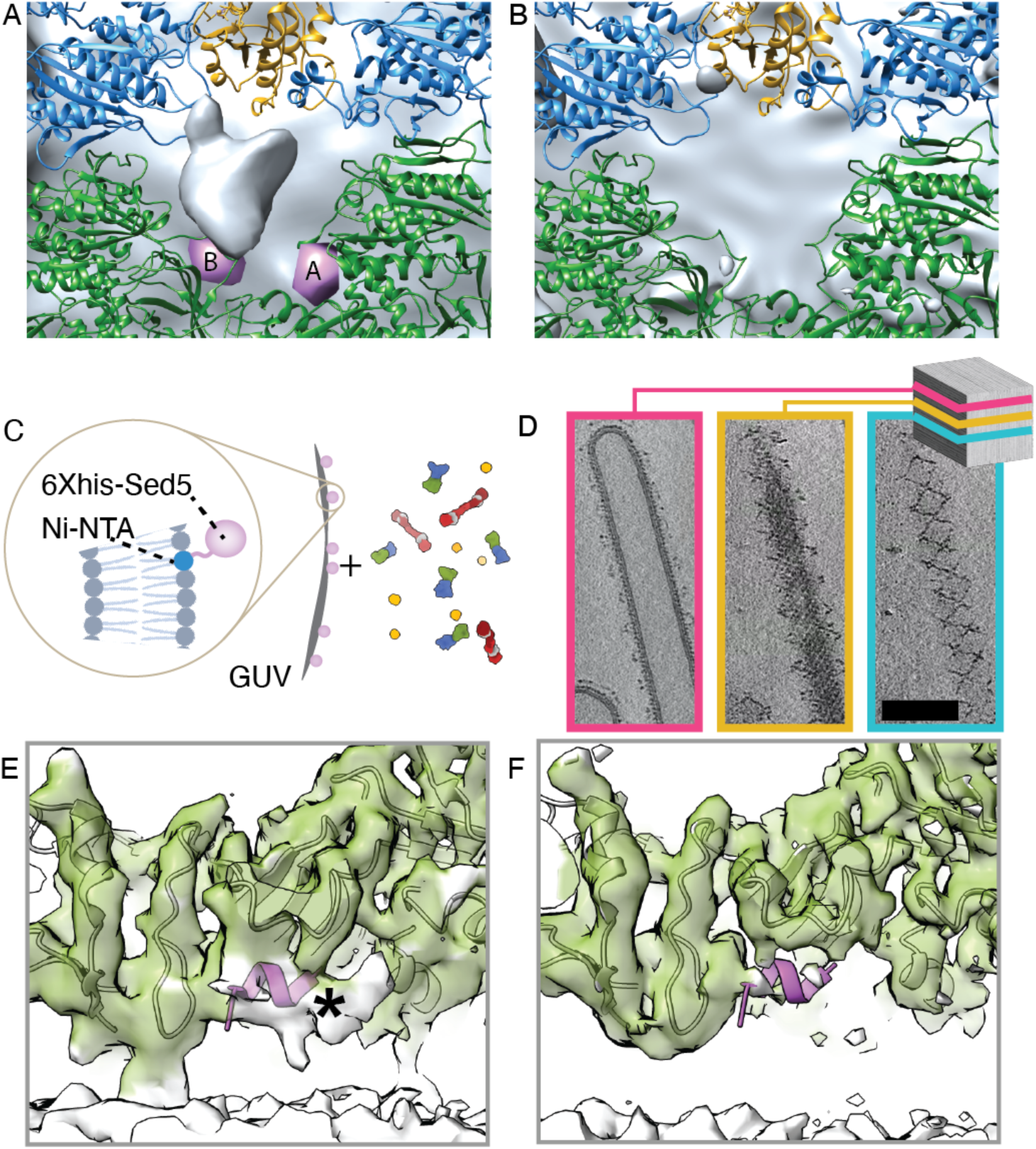
(A) Difference map between the subtomogram average of the inner coat on microsomes and a 14 Å low-pass filtered volume representation of the fitted model of the inner coat from cargo-less GUVs (PDB: 8BSH) with Sar1 in yellow, Sec23 in blue, and Sec24 in green. The Sec24 A- and B- cargo-binding sites are represented with pink blobs generated from bound cargo peptides described in previous X-ray crystallography studies (PDB: 1PD0, 1PCX). (B) as in (A) but using the STA map previously obtained from GUVs (EMD-15949), and low-pass filtered to 14 Å. (C) Schematic of the in vitro reconstitution of COPII budding from Sed5-enriched GUVs, the inset highlights details of His_6_-tagged Sed5 associating to the Ni- NTA tagged lipids on the GUVs. (D) XY slices through a representative tomogram of Sed5- enriched GUV budding reactions at different Z heights displaying the coated tube morphology (pink, z=162), and the inner (yellow, z=137) and outer (blue, z=128) coat arrangements. Scale bar = 100 nm. (E) Detail of the STA map of Sed5-bound inner coat showing the region around the Sec24 B-site. Density closer than 3.5 Å to the fitted model of the inner coat (PDB: 8BSH) is shown in green whilst white density corresponds to regions of the map that are not explained by the fitted model. The model of a Sec24-bound peptide from Bet1 cargo (PDB: 1PCX), which contains the same B-site binding motif as Sed5 (LxxLE), is also fitted to highlight the location of the B-site and is shown in purple. White density in correspondence of this peptide is marked with an asterisk. (F) As in (E) but displaying the map obtained from cargo-less GUV reconstitution (EMDB-15949).

To further support the hypothesis that the extra density we see is bona fide cargo, we analysed the composition of microsomal membranes by mass-spectrometry. We detected many known Sec24 cargo proteins among the most abundant constituents of these membranes. These include export receptors (Erv25, Erv29, Erv46, Erv41, Emp24, Svp26, several Erp proteins), components of the Golgi SNARE complex (Sec22, Sed5, Bet1, Bos1), HDEL receptor (Erd2), and other Sec24 interactors (Yor1, Prm8, Shr3). We also detected a high number of proteins that are localised to other compartments such as the plasma membrane, Golgi, and endosomal network. While these could come from contaminant membranes in the microsome prep, we assume a subset belong to a newly synthesised pool located in the ER and ready to enter COPII vesicles. Cargo proteins and recycling receptors have been previously shown to be enriched in COPII-coated vesicles over prevailing concentration in microsomal membranes^26,27^.

Due to the presence of a wide range of structurally diverse cargoes on the microsomal membranes, it was not possible to resolve the bound protein density to anything other than a shapeless ‘blob’ (Figure 3A). It is likely that the cargoes bound to the inner coat within the lattice are either small, flexible, or both, as cargoes with bulky cytosolic domains would be sterically prevented from binding in the 50Å-wide space between neighbouring heterotrimers (Figure 3A). Whilst we expect different subsets of Sec24 molecules to be bound to cargos of different sizes, or not at all, we were unable to reproducibly differentiate between them using 3D classification. This is likely due to the high amount of compositional and conformational heterogeneity of the cargo molecules, and the fact that different sites on Sec24 may be bound substoichiometrically to different cargoes.

To further test whether inner coat lattice formation is compatible with cargo binding, we reconstituted COPII budding using GUVs whose surface was enriched with the cytosolic domain of a small cargo protein, Sed5 from *S. cerevisiae*. Sed5 is a SNARE protein that acts at the cis-Golgi in complex with other SNARES^28^. Sed5 contains two known Sec24 binding motifs specific for the A- and B-sites (YNNSNPF and LMLME, respectively)^25^. Biochemical studies have shown that the YNNSNPF motif on Sed5 is occluded in the monomeric state but becomes exposed in the context of the SNARE complex^25^, raising interesting questions about regulation of its transport. An analogous conformational switch has been reported for other SNARE proteins of the syntaxin family^29,30^. AlphaFold predictions of Sed5 structure (PDB ID: AF-Q01590-F1) suggest that both Sec24 binding peptides are found in a highly flexible region characterised by very low confidence scores, allowing Sed5 to bind in the small space between inner coat units (Extended Data Fig. 2A).

First, we purified the Sed5 cytosolic domain (residues 1 to 319) to high purity and homogeneity (Extended Data Fig. 2B). We enriched the surface of the GUVs with Sed5 by the association of Ni-NTA tagged lipids in the GUVs to a C-terminal His_6_ tag in the purified Sed5, cloned in place of the transmembrane domain (Figure 3C, inset). We verified the successful association of Sed5 to the membrane by liposome flotation assays (Extended Data Fig. 2B). We then carried out COPII budding reconstitution *in vitro* using Sed5-enriched GUVs (Figure 3C). Imaging these budding reactions using cryo-electron tomography revealed that COPII primarily forms tubes (88.8 % of all coated membranes) (Figure 3D), similarly to previous studies with cargo-less GUVs^15^. The inner and outer coat lattices were clearly visible on these tubes (Figure 3D).

To establish whether Sed5 was bound within the inner coat lattice, we carried out STA to generate a high-resolution (4.1 Å) structure of the inner coat lattice (Extended Data Fig. 3A-C). The Sed5-bound map was essentially identical to previous structures lacking cargo, but crucially, we saw unambiguous protein density in one of the known Sed5 binding pockets in correspondence to the B-site (Figure 3E and F). We were unable to resolve any further Sed5 protein density outside of the known binding pocket on Sec24. This is unsurprising given that the Sec24 binding motifs on Sed5 were predicted to be in a highly flexible and disordered region (Extended Data Fig. 2A). The A site appears unoccupied (Extended Data Fig. 3), consistent with previous studies that showed that the A-site specific YNNSNP peptide is occluded in the monomeric form of Sed5. This peptide only becomes exposed when Sed5 forms SNARE complexes with its partners, and thus monomeric Sed5 is expected to bind the B-site^25^. We have confirmed that Sed5, as presumably other small and flexible cargo proteins, can bind to the inner coat without disrupting the lattice. The lattice disruption we observe on vesicles derived from native microsomes is therefore probably due to the presence of more bulky proteins. We note that, given the purity of the Sed5 prep (Extended Data Fig. 3) and the absence of any additional differences with the cargo-less budding reaction, it is highly likely that the extra density we see corresponds to Sed5.

### The COPII Outer Coat is Heterogeneous on Vesicles

The Sec13-31 outer coat layer was clearly visible on microsome-derived COPII-coated vesicles (Figure 1B). Manual inspection of denoised tomograms revealed that the outer coat was generally arranged in cage-like structures, with “rods” of Sec13-31 acting as edge elements. Multiple arrangements of these Sec13-31 rods were observed, as shown in some example cages depicted in Figure 4A. In many instances, four rods converge to form vertices through the interaction of Sec31 N-terminal beta-propeller domains, in the canonical manner previously described for *in vitro* assembled protein-only cages, and reconstituted tubules^10,11,15^ (Figure 4A bottom panels, red spheres). We also observed rods where one or both of the Sec31 N-terminal domains bind to the Sec31 dimerisation domain (Figure 4A bottom panels, dotted lines). We previously described a similar interaction on tubules, and proposed that it stabilises the outer coat when neighbouring patches are ‘out of phase’ with respect to one another and vertices cannot form^15^. Finally, we observe an entirely novel interaction where 5 rods converge to form vertices (Figure 4A bottom panels, blue spheres). The edge elements in the cages outline triangular, rhomboidal, as well as pentameric faces (Figure 4A). These were previously described in in vitro-assembled protein-only cages, where regular cuboctahedral and icosidodecahedral structures were formed in high-salt^10,11^, However, in our physiological buffer conditions and while coating native membranes, no such global symmetry was detected.

**Figure 4.**
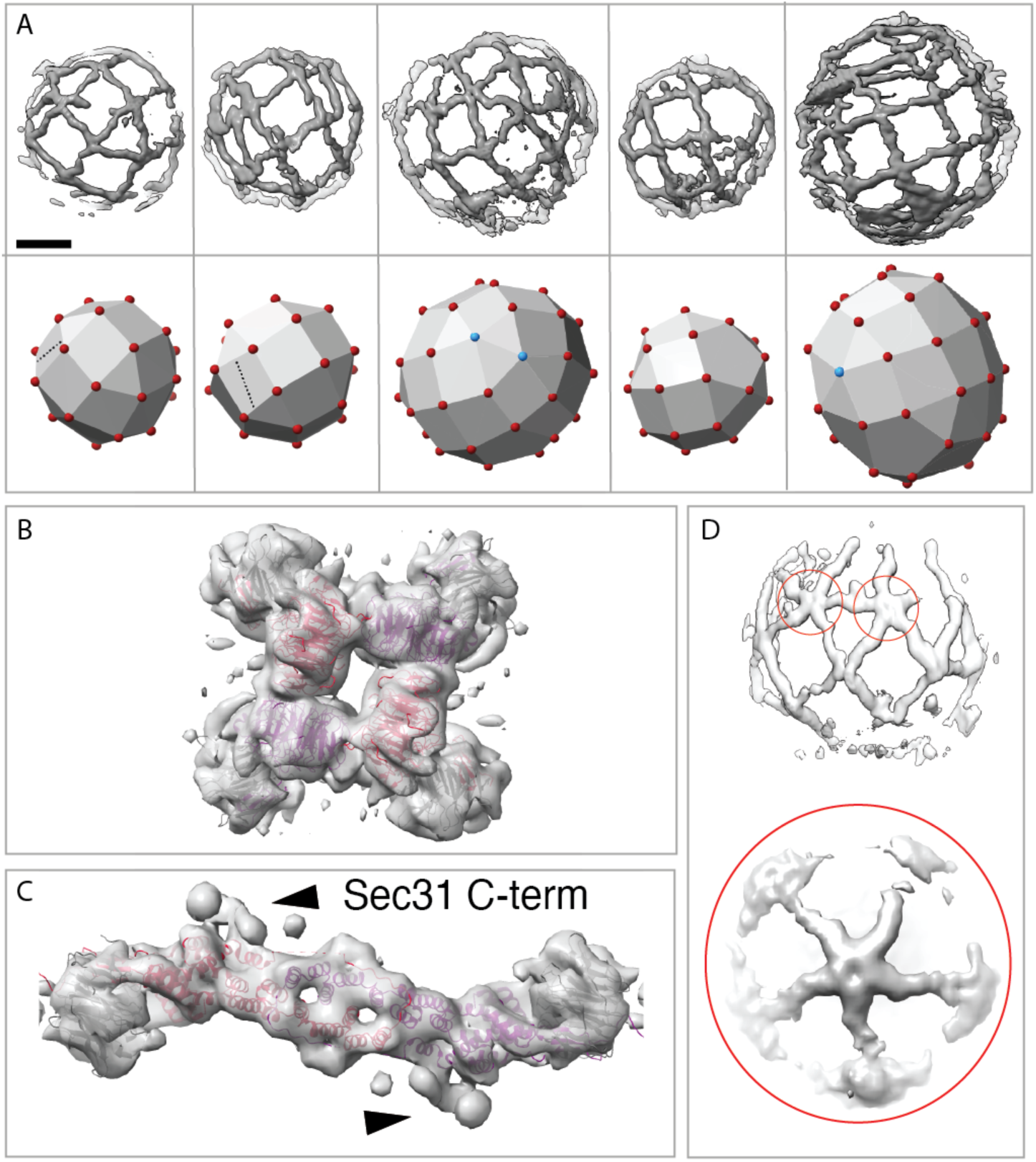
(A) COPII outer coat arrangement on vesicles. Top panel: outer coat as visible in denoised tomograms. Tomograms were masked for visual clarity. Bottom panel: schematic representation of the cage architecture seen in the top panels. Red and cyan spheres indicate vertices with 4 or 5 rods convergingrespectively, dotted lines indicate rods forming non-vertex interactions. Scale bar = 30 nm. (B) STA map of the outer coat vertex on vesicles from microsomes at 11.4 Å, with four copies of the atomic model of the Sec13-Sec31 ‘vertex element’ fitted (PDB: 2PM9) with Sec31 in red and purple, Sec13 in grey (C) STA map of the outer coat rod on vesicles at a resolution of 11.5 Å, with the atomic model of the Sec13- Sec31 ‘edge’ element fitted (PDB: 2PM6). Colour code as in (B). (D) Structure of vertices formed by convergence of 5 rods. Top panel, an example tomogram density depicting two 5- way vertices (red circles). Bottom panel, the STA average from 460 manually picked 5-way vertex particles.

The structures of the canonical outer coat vertex and Sec13-Sec31 rods were resolved by STA to 11-12 Å for both the microsome-derived and the Sed5-enriched GUV samples. This resolution allowed unambiguous rigid-body fitting using previously determined atomic models (PDB: 2PM9 and 2PM6) (Figure 4B-C, and Extended Data Fig. 4). For both vertices and rods, the Sed5-GUV and microsome-derived maps are very similar (Extended Data Fig. 4). Previously, we showed that the Sec31 C-terminal domain binds to the dimerization domain of another Sec31 to form an ‘elbow’ (Figure 4C, black arrowheads) and hypothesized that this interaction is important to stabilise the coat. However, the microsome-derived structure contained a stronger and better-defined density for the C-terminal domain of Sec31 (Extended Data Fig. 4E-F). Taken together, this suggests a more prominent role for this stabilising interaction in the context of the widely varying assembly seen on the spherical vesicles derived from microsomes compared to the GUV-derived tubules. We also solved the structure of the vertex formed by convergence of five rods (Figure 4D). Although the resolution is low due to the limited number of particles, the shape and size of Sec13-Sec31 rods is clearly recognisable, indicating that these vertices are formed by interaction of five Sec31 beta-propeller domains.

The refined positions and orientations of vertex and edge elements obtained by STA allow us to make quantitative analysis of cage architecture. When we plot the positions of neighbouring vertices for each vertex (Figure 5A), we notice that neighbours cluster at expected positions, being roughly 300 Å apart and forming an ‘X’ shape. However, their diffuse ‘cloud’ distribution clearly shows a wide range of possible angles formed at vertices, both tangential (Figure 5A, left panel), as well as normal to the membrane (right panel). We measured the average angle below each vertex (alpha). We found it can assume values between 120 and 180 degrees, and that these change continuously rather than clustering in ‘preferred’ angles, as also seen in the neighbour plot. For each vertex, we plot alpha angle versus vesicle diameter and we find a strong positive correlation (Figure 5B), indicating that outer coat structure is related to membrane curvature.

**Figure 5.**
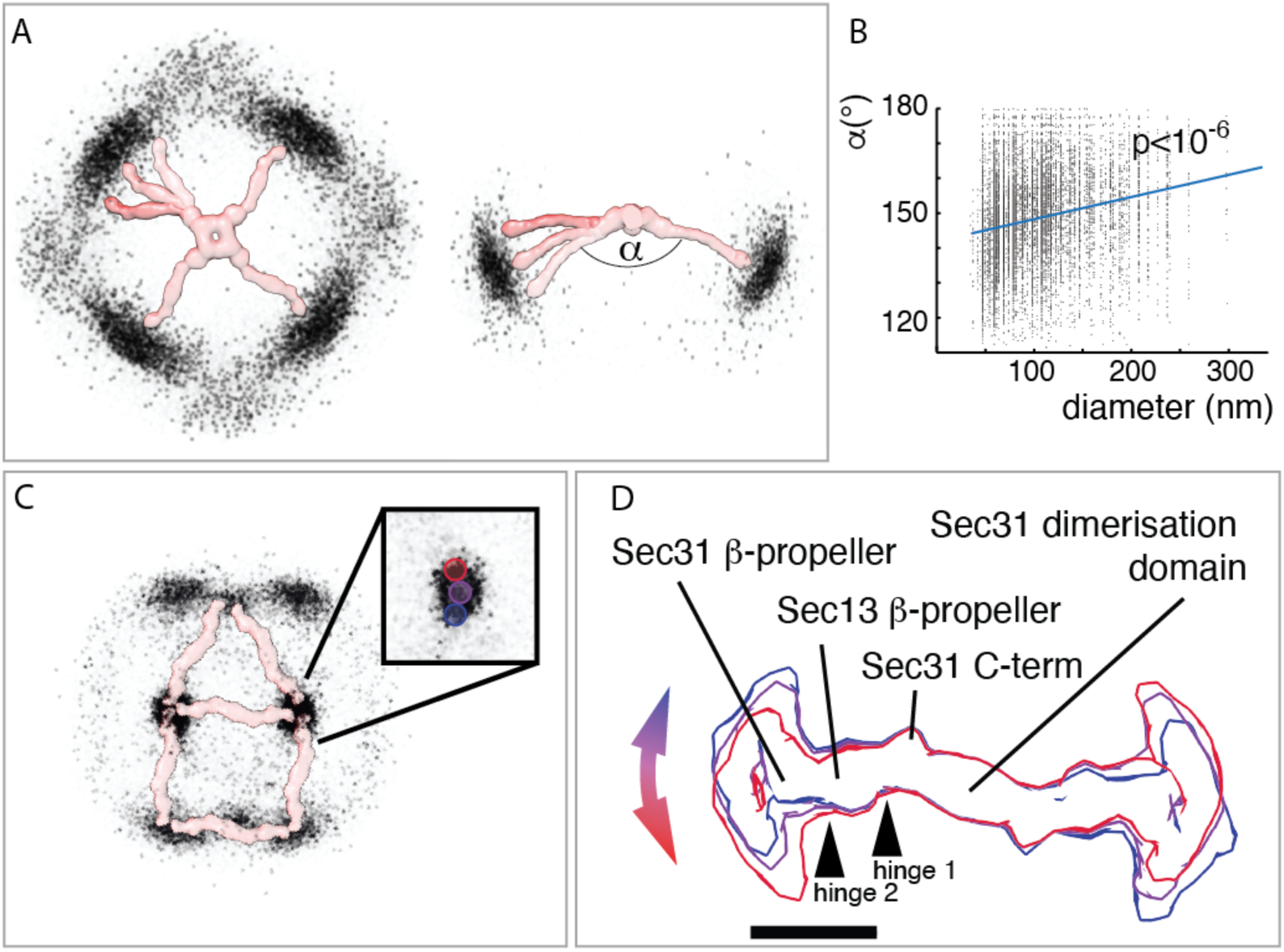
(A) The position of neighbouring vertices surrounding all vertices is plotted: each black dot corresponds to the position of one neighbour. Filtered density maps for vertices and rods are manually overlaid to aid visual interpretation. Left and right panels: view from the top and side respectively. (B) The average angle below each vertex (as shown in panel (A)) is plotted against the corresponding vesicle diameter, together with correlation trendline and p value (2-tailed t-test, p=5.45E-07). (C) The position of vertices surrounding all rods is plotted: each black dot corresponds to the position of one vertex. The pattern appearing from clustering of nearest vertices corresponds to most rods being arranged in rhomboids and triangles (overlaid rod density). Inset: zoom in one cluster of neighbour positions, showing masks used to select particle classes (D) Variation of rod structures. Rods were selected according to the nearest vertices falling within regions defined by the red, purple and blue masks shown in the inset in (C) and were reconstructed as different classes. The resultant structures are overlayed and show movement around two major hinge regions. Scale bar = 10 nm.

Analysis of the arrangement of the outer coat revealed further unexpected heterogeneity. By plotting the positions of the nearest vertex neighbours for all rods (Figure 5C), a general rhomboidal and triangular pattern emerged conveying the expected outer coat arrangement of rods with a vertex at each extremity. However, the points representing the positions of the vertices relative to the centre of each rod did not form a sharp peak. Instead, the distribution of the distance of the vertices relative to the rods was broad, which suggested that the rods are not rigid. To investigate this further, we defined distinct classes of rods based on their distance to their nearest vertex from within the ‘cloud’ of points (blue, purple, and red masks in Figure 5C inset). We reconstructed the corresponding classes of rods, generating three different maps which demonstrated variation in rod structure (Figure 5D). Specifically, this analysis revealed high mobility around two previously unidentified hinge regions located near the Sec13- and Sec31- β-propellers (Figure 5D, Supplementary Movie 1).

In summary, the variety of arrangements presented here suggests that the outer coat is highly morphologically heterogenous on native vesicles, in stark contrast to the more regular outer coat morphology seen on tubes and on *in vitro* assembled membrane-less polyhedral cages.

### Relationship between inner and outer coat

Finally, we analysed the relationship between the inner and outer coat on vesicles. Previous studies established that Sec31 binds to Sec23/Sar1 via a 300 aminoacid long flexible linker^25,31^.

On coated tubules, we had previously found that this flexible linker allows for the outer coat layers to float on top of the inner layer. This allows for the two lattices to adapt to a continuous range of membrane curvatures^15^. We also found that whilst there is no fixed alignment between the inner and outer coat lattices on tubules, both layers are rotationally aligned, as they both follow a helical pattern around the tubules.

To assess whether the Sec31 flexible linker allows some degree of movement of the outer coat bound to the inner coat on native membranes, we plotted the position of inner coat subunits neighbouring each outer coat vertex (Extended Data Fig. 5A). The neighbour positions cluster below vertices at the expected distance, but assume a very broad distribution that does not show any pattern. This suggest there is no fixed alignment between the inner and outer coat lattices, similar to what we had described for coated tubules^15^.

To see whether the Sec31 flexible linker allows for rotation of outer coat with respect to the inner coat, or whether the two lattice layers are locally rotationally aligned as seen on tubules^11^, we selected a subset of vertices that all had a neighbouring inner coat subunit within a limited region (yellow mask in Extended Data Fig. 5A).

If the two layers were rotationally aligned, we would expect that the average of a subset of vertices with fixed translational relationship to neighbouring inner coat subunits would show clear density for both layers. Averaging the selected vertex subset produced a map where no discernible inner coat structure was visible below the vertex, indicating the two layers are both translationally and rotationally unrelated (Extended Data Fig. 5B). This suggest the flexible linker allows for full translational and rotational freedom, and that there are no other factors binding the two layers together in our in vitro reconstitution.

## Discussion

We reconstituted COPII budding *in vitro* using *S. cerevisiae* microsomes as native membrane sources. Microsomes are cell-derived membranes, purified by sucrose gradient centrifugation, which largely comprise of the ER. Therefore, microsomes resemble the ER in their lipid composition, heterogeneity, and importantly, the presence of transmembrane and lumenal cargo proteins. In striking comparison to the COPII-coated tubules generated from GUVs, microsome-derived membrane carriers are mostly pseudo-spherical. The overall morphology and appearance of the coat reconstituted on microsomes is very reminiscent of the structures seen in situ on cryo-FIB/SEM data obtained from Chlamydomonas reinhardtii^21^ and yeast (EMPIAR-11462^32^).

The vast majority of vesicles were detached from the donor membranes, with only a handful of instances of constricted necks. Here, we use a non-hydrolysable GTP analogue, suggesting GTP hydrolysis is not required for scission in our system, consistent with previous studies^33^. While we do not perform any centrifugation or mechanical perturbation aside from gentle pipetting and blotting to prepare our samples, we can’t exclude scission is triggered by non-physiological mechanisms. However, similar experiments performed on GUVs with COPII interface mutants resulted in coated spherical profiles that remained linked by constricted necks (like beads on a string)^15^, suggesting that scission might depend on factors present within the microsome membrane.

Subtomogram analysis of the coat on spherical vesicles showed that the inner coat Sar1-Sec23-Sec24 heterotrimers assemble into small patches of lattice, in contrast to the continuous lattice found on tubules. The arrangements of neighbouring subunits in these small patches and in the extended lattice found on regular tubules are highly similar. Cargo protein density was detected within the inner coat lattice of microsome-derived vesicular carriers, indicating that small and/or flexible cargo cytosolic domains can be accommodated within the tightly packed inner coat. This was further confirmed by the reconstitution of the flexible Sed5 cytosolic domain onto GUVs, as COPII budding leads to formation of highly ordered tubules where a short Sed5 peptide can be detected bound to Sec24 B-site. We do not see any significant density for the Sed5 peptide bound to the A-site, consistent with previous descriptions of occlusion of the A-site interaction motif in the monomeric SNARE^25^ (Extended Data Fig. 3D-E).

Subtomogram analysis of the coat assembled around spherical vesicles also revealed a highly variable and flexible outer coat cage consisting of Sec13-31 rods assembling with many different geometries. Most rods converge through interactions between Sec31 N-termini to form canonical 4-way vertices, but we also detected T-junctions with other rod’s dimerization domain, as well as pentameric vertices. Outer coat cages have elements of icosidodecahedral and cuboctahedral geometry, such as triangular, rhomboidal and pentameric faces, reminiscent of membrane-less in vitro assembled cages. However, individual cages are distinct and no overall symmetry is present. The rods themselves are highly flexible, with two major hinge points around the Sec13 β-propeller.

The interaction between the inner and outer coat layers, known to be mediated by a disordered region of Sec31, is also variable with local inner and outer coat lattices being translationally and rotationally not aligned.

Overall, our findings that COPII morphology differs between microsomes and naked GUVs, in combination with our previous finding that the regions responsible for outer coat assembly are not necessary for budding^15^, challenge the idea that the outer coat cage assembly is the main driver of membrane curvature. We propose a new model for generation of membrane curvature by the COPII coat (Figure 6), where vesicle shape is mostly determined by inner coat assembly. According to this model, the extent of inner coat lattice polymerisation drives membrane curvature. In the extreme case of undisturbed lattice assembly, as obtained with GUVs *in vitro* where GTP hydrolysis is inhibited, no bulky proteins are present, and membrane sources are abundant, coated tubes are formed. In native conditions, where bulky proteins are present to disrupt inner coat lattice assembly, small patches of randomly oriented inner coat lattice will lead to formation of pseudo-spherical vesicles. In this scenario, the outer coat’s ability to adapt to a continuous and varied range of growing curvature ensures effective binding and assembly of cages, which stabilise the coated vesicle. Of note, *in vitro* reconstitutions from GUVs using COPII mutants with weakened inner coat lattice interfaces also led to the formation of spherical profiles^15^, supporting the proposed model.

**Figure 6:**
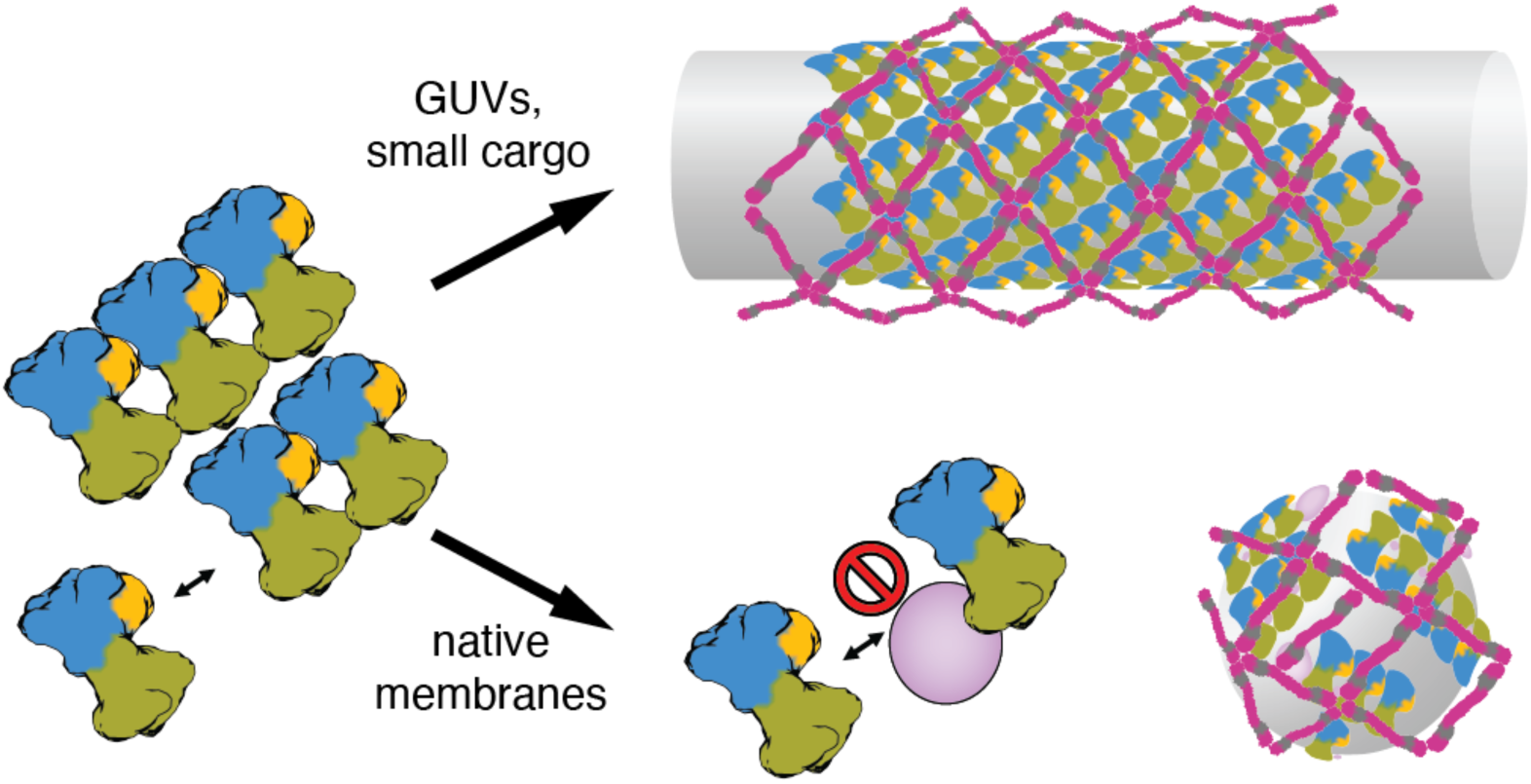
Schematic describing the proposed model of COPII budding, where inner coat lattice assembly drives curvature of the membrane. If undisturbed, this leads to an extended pseudo-helical lattice and formation of coated tubes (top). In native conditions, where bulky protein (pink) are present, extensive assembly of inner coat is not possible and small patches randomly orient to generate near-spherical membranes

## Acknowledgements

We thank Natasha Lukoyanova and Shu Chen at the ISMB Birkbeck Cryo-EM Lab, Zhengyi Yang and Wim Hagen at the EMBL Imaging Centre in Heidelberg, and eBIC for cryo-tomography data collection, David Houldershaw at Birkbeck College for computational support, and Katie Downes at the Crick for help with the figures. We thank lab members Sander van der Verren and Katie Downes for comments on the paper.

We acknowledge the access and services provided by the Imaging Centre at the European Molecular Biology Laboratory (EMBL IC), generously supported by the Boehringer Ingelheim Foundation. We acknowledge the ISMB EM facility (Birkbeck College, University of London), supported by the Wellcome Trust (202679/Z/16/Z and 206166/Z/17/Z). We thank Farida Begum for help with mass spectrometry analysis, which was performed at the Biological Mass Spectrometry and Proteomics Facility of the MRC-LMB.

This work was supported by grants to GZ from the European Research Council (ERC-StG-2019 grant 852915) and the BBSRC (BBSRC grant BB/T002670/1) to G.Z. Work in the group of EAM was supported by the Medical Research Council, as part of United Kingdom Research and Innovation (also known as UK Research and Innovation) [MC_UP_1201/10]. For the purpose of open access, the MRC Laboratory of Molecular Biology has applied a CC BY public copyright licence to any Author Accepted Manuscript version arising.

## Author Contributions

Conceptualisation: G.Z.; Funding acquisition: G.Z.; Sample preparation for cryo-EM: E.P.; Cryo-electron tomography data collection: E.P..; Cryo-electron tomography and STA data processing: E.P., G.Z.; Microsome preparation and mass spectrometry: E.A.M. Writing: (original draft) E.P., G.Z., (revisions) E.P., E.A.M., G.Z.

## Competing Interests

The authors declare that no competing interests exist.

## Materials and Methods

### Cloning

Sed5 (UniProt: Q01590) (residues 1-319, truncating the transmembrane helix) was cloned from the *Saccharomyces cerevisiae* S288c genome into a pETM-11 expression vector linearised at the XhoI and XbaI restriction sites using In-Fusion (Takara) technology. A flexible triple-glycine linker was added between the C-terminal residue (319) of Sed5 and a His_6_ tag. The primers are used were:

Fo: 5’- GGTGTCCTCCTCCTCTATTACTCTTTATCCTGTCGAAG -3’

Re: 5’- GAGGAGGAGGACACCACCACCACCACCAC -3’

Sec23/24, Sec13/31, and Sar1 constructs previously described in Hutchings *et al.* (2021)^15^ were used here.

### Protein Expression and Purification

The Sed5 pETM-11 vector was transformed into *Escherichia coli* (BL21) cells via heat shock. Cells were cultured at 37 °C with 220 rpm shaking in 2 L of LB media supplemented with kanamycin. When cultures reached an optical density between 0.7-1, 0.2mM IPTG was added, and the incubation temperature reduced to 16 °C. Culture pellets were harvested after approximately 16 hours by centrifugation and flash frozen in liquid nitrogen before storage at −80 °C.

Sed5 pellets from 2 L culture were thawed and resuspended in 20 mL Ni-A buffer (50 mM Tris (pH 8), 500 mM KCl, 0.1 % TWEEN 20 (v/v), 10 mM imidazole, 1 mM DTT) supplemented with 1 complete protease inhibitor tablet (Roche). 40 mg/mL lysozyme was added to the pellets, which were then stirred on ice for 20 minutes. Cells in the pellets were lysed using a cell disruptor. Unbroken cells were removed by ultracentrifugation at 20200 *xg* for 25 minutes. The supernatant was loaded onto a Ni-NTA 5 mL His-trap column (G.E. Biosciences) equilibrated with Ni-A buffer and washed with 5 column volumes of Ni-A buffer. Sed5 was eluted from the column via applying a linear gradient of Ni-B buffer (50 mM Tris (pH 8), 500 mM KCl, 0.1 % TWEEN 20 (v/v), 500 mM imidazole, 1 mM DTT). Fractions were analysed by SDS-PAGE and those containing Sed5 were pooled before 10x dilution in Q-A buffer (20mM Tris (pH 8.0), 0.1 % TWEEN-20 (v/v), 10 % glycerol (v/v), 1 mM DTT). Sed5 was loaded onto a 5 mL HiTrap Q column (G.E. Biosciences) equilibrated with Q-A buffer. The column was washed with 2 column volumes of Q-A buffer, and 2 column volumes of a mixture of 90 % Q-A buffer and 10 % Q-B buffer (20mM Tris (pH 8.0), 0.1 % TWEEN-20 (v/v), 10 % glycerol (v/v), 1 mM DTT, 1 M KCl). Sed5 was eluted with a linear gradient of Q-B buffer. Fractions were analysed by SDS-PAGE and those containing Sed5 were pooled and concentrated using a protein concentrator with a 10 kDa molecular weight cut off to a final concentration of 0.5 mg/mL. Sed5 was separated into 100 μL aliquots and flash frozen.

The final step of Sed5 purification was carried out on the day of use. 1 aliquot of Sed5 was thawed before injection onto a Superdex 200 Increase 3.2/300 column equilibrated with HKM buffer (20 mM HEPES, 50 mM KOAc and 1.2 mM MgCl2, pH 6.8). Fractions containing Sed5 were identified by SDS-PAGE and pooled together.

The purified protein was confirmed as Sed5 by analysis with SDS-PAGE combined with gel sequencing by mass spectrometry at the Mass Spectrometry and Proteomics Facility at the University of St. Andrews.

Sec23/24, Sec13/31, and Sar1 were expressed and purified as described previously from SF9 and e.Coli cells, including the steps to cleave the His_6_ tags in Sec23/24 and Sec13/31^15^.

### Liposome Flotation Assays

Liposomes were generated as previously described^34^ using the ‘Major-Minor’ lipid mixture: 49 mol% phosphatidylcholine, 20 mol% phosphatidylethanolamine, 8 mol% phosphatidylerine, 5 mol% phosphatidic acid, 9 mol% phosphatidylinositol, 2.2 mol% phosphatidylinositol-4-phosphate, 0.8 mol% phosphatidylinositol-4,5-bisphosphate, 2 mol% cytidine-diphosphate-diacylglycerol, supplemented with 2 mol% TexasRed- phosphatidylethanolamine, 2 mol% Ni-NTA tagged lipids (18:1 DGS-NTA(Ni)), and 20% (w/w) ergosterol.

Liposomes were pre-mixed with the Sed5, and floatation assay experiments were performed without and with the addition of COPII components: 1 μM Sar1, 180 nM Sec23/24, 173 μM Sec13/31, 360 nM Sed5 with 1 mM GMP-PNP (Sigma-Aldrich), 2.5 mM EDTA (pH 8.0). All floatation assays contained 0.27 mM liposomes in a total volume of 75 μL. Liposome floatation reactions were mixed with 250 μL 1.2 M sucrose in HKM buffer in an ultracentrifuge tube. 320 μL 0.75 M sucrose in HKM was gently layered on top. A final layer of 20 μL HKM was then layered on top of the sucrose solutions. Ultracentrifuge tubes were loaded into a SW-55 Ti ultracentrifuge rotor before spinning at 280k *xg* at 4 °C for at least 16 hours. The top 20 μL of the sucrose gradient was carefully extracted before analysis via SDS-PAGE with silver staining.

### Budding Reactions

Purified microsomes from *S. cerevisiae* were prepared as described previously^31^. 1.5 mg of microsomes were washed three times carrying out the following steps: resuspending the microsomes in 1 mL B88 buffer (20 mM HEPES (pH 6.8), 150 mM KOAc, 250 mM sorbitol, 5 mM Mg(OAc)2), pelleting membranes by centrifugation on a chilled benchtop centrifuge at 20,000 *xg* for 2 minutes, removing the supernatant, and resuspending the pellet in 50 μL B88 buffer. After washing, the pellets were diluted a further 8x, and chilled on ice before use in budding reactions.

Budding reactions in microsomes were prepared by incubating 1 μM Sar1, 180 nM Sec23/24, 173 μM Sec13/31 with 1 mM GMP-PNP (Sigma-Aldrich), 2.5 mM EDTA (pH 8.0) and 10 % microsomes (v/v).

GUVs were prepared by electroformation^35^ from 10 mg/mL of a major-minor lipid mixture with 2 mol% Ni-NTA tagged lipids (described in the liposome floatation assay section) in a 2:1 chloroform:methanol solvent mixture, as described previously^14,36^. The lipid mixture was spread over two Indium Tin Oxide-coated glass slides. 300 mM sucrose was suspended in a silicon O-ring between these glass slides and GUVs were generated using a NanIon Vesicle Prep Pro. GUVs in the sucrose solution were added to 500 μL of 300 mM glucose and left to sediment overnight at 4 °C. The supernatant was discarded, leaving a 50 μL pellet of GUVs.

Budding reactions in GUVs with Sed5 were prepared by incubating 1 μM Sar1, 180 nM Sec23/24, 173 μM Sec13/31, 360 nM Sed5 with 1 mM GMP-PNP (Sigma-Aldrich), 2.5 mM EDTA (pH 8.0) and 10 % GUVs (v/v). GUVs were pre-mixed with the Sed5 prior to addition to the COPII components. Budding reactions were incubated for at least 30 minutes before vitrification for cryo-electron tomography.

### Cryo-Electron Tomography Sample Preparation

5 nm BSA-blocked gold nanoparticles (BBI Solutions) were added to the budding reactions at a concentration of 10 % (v/v). 4 μL of budding reactions from GUVs or microsomes was added to glow-discharged Lacey Carbon Films on 300 Mesh Copper Grids (Agar Scientific), incubated for 60 seconds, before back-blotting on a Leica-GP2 plunge freezer in 95 % humidity and a 4 second blotting time. Vitrified grids were stored in liquid nitrogen prior to data collection.

### Cryo-Electron Tomography Data Collection

Budding reactions with microsomes were imaged using cryo-electron tomography at the EMBL Imaging Center in Heidelberg on a Titan Krios microscope (Thermo Scientific) operated at 300 kV. The microscope was equipped with a SelectrisX energy filter (Thermo Scientific) and a Falcon 4 detector (Thermo Scientific) in counting mode. Pixel size was 1.526 Å and tilt series were taken with a defocus range of −3 μm to −5 μm. Tilt series were aquired using a dose-symmetric tilt scheme^37^ over a total exposure of 140 e^-^ Å^-2^ with tilt angles ranging between −60 ° to +60 ° at a 3 ° increment. Data collection was controlled using SerialEM^38^ and implementing PACE-tomo^39^. 765 high-quality tilt series were collected, yielding the same number of tomograms.

Budding reactions with GUVs and Sed5 were imaged using cryo-electron tomography at the EMBL Imaging Center in Heidelberg over two sessions of data collection on a Titan Krios microscope operated at 300 kV. The microscope was equipped with a K3 (Gatan) detector and energy filter. The first session was collected in super-resolution mode and the second session was collected in counting mode. Pixel size was 1.33 Å and tilt series were taken with a defocus range of −1.5 μm to −3.5 μm. Tilt series were aquired using a dose-symmetric tilt scheme^37^ over a total exposure of 142 e^-^ Å^-2^ with tilt angles ranging between −60 ° to +60 ° at a 3 ° increment. Data collection was controlled using SerialEM. 326 high-quality tilt series were collected, yielding the same number of tomograms.

Grids were screened and optimised at the ISMB EM Facility at Birkbeck College.

### Cryo-Electron Tomography Data Processing

The microsomes dataset was processed using an Alpha-phase development version of RELION 5.0 (4.1-alpha-1-commit-d2053c) (manuscript in preparation). Initially, .mdoc files were renamed TS_[number]-style to ensure compatibility with RELION to Dynamo (and vice versa) scripts used later in the processing workflow. Raw data then was imported into RELION 5.0. Individual tilt movies were motion corrected and averaged using whole frame alignment in the RELION implementation of MotionCor2^40,41^. CTF estimation was carried out using CTFFIND-4.1^42^ with a defocus range of −25000 to −55000 Å and a maximum CTF resolution of 20 Å. Tilt series were manually inspected and poor tilt images were removed using a Napari plug-in (https://github.com/napari/napari/blob/main/CITATION.cff) provided as part of the Exclude tilt-images job-type in RELION 5.0. Tilt series were automatically aligned using the IMOD wrapper for fiducial-based alignment in RELION with a fiducial diameter of 8 nm. Tomograms were reconstructed in RELION at a pixel size of 12.208 Å for visual inspection and particle picking. Tomograms were denoised and missing wedge-corrected using IsoNet for use in manual particle picking^43^. We also generated 8x binned CTF-corrected tomograms for use in PyTOM template matching^44^, using IMOD’s *etomo*^45^ function on the IMOD metadata generated by the Align tilt-series job-type in RELION.

The GUVs-Sed5 dataset was processed using the RELION4_Tomo_Robot (https://github.com/EuanPyle/relion4_tomo_robot/blob/master/CITATION.cff). Individual tilt movies were motion corrected and averaged using whole frame alignment with MotionCor2^40^. Movies collected in super resolution mode were binned by 2x during motion correction. Tilt series were created from individual tilt images using IMOD’s *newstack* function. Tilt series were manually inspected using IMOD’s *3dmod* visualisation function and bad tilts were removed using IMOD’s *excludeviews* function. Tilt series were automatically aligned using Dynamo’s automated fiducial-based alignment in the RELION4_Tomo_Robot’s ‘fast_mode’ with a fiducial diameter of 5 nm^46^. CTF estimation was carried out using CTFFIND-4.1^42^. The dataset was then imported into RELION 4.0^16^. Tomograms were reconstructed in RELION at a pixel size of 10.64 Å for visual inspection and particle picking. Tomograms were denoised and missing wedge-corrected using IsoNet for visual inspection^43^.

### Subtomogram Averaging

#### Microsomes Dataset

##### Inner coat

The surface of vesicles in IsoNet-denoised tomograms was defined and segmented using the Pick Particle plug-in in Chimera as described previously^47,48^. The coordinates of the vesicle surface were used to mask the tomograms to enable manual particle picking in UCSF Chimera which were assigned Euler angles normal to the membrane. 3579 particles were extracted in 48 voxel boxes from RELION-reconstructed (non-denoised) tomograms at a voxel size of 9.156 Å. Particles were assigned random in-plane rotation angles and were averaged to create a reference using Dynamo^46^ with 4697 particles. Particles were then aligned and averaged in Dynamo with the following conditions: a cone range of 10° was applied whilst 360° of in-plane rotation was allowed; particle translation was limited to 1 voxel in all directions due to the accuracy of the coordinates of the manually picked particles; a C2 symmetry was applied due to the pseudo-symmetry of the inner coat at low resolution; a mask covering the area of one inner coat subunit was applied; alignment was carried out for 100 iterations. The resulting Dynamo table was converted to a .star file using *dynamo2relion* (https://github.com/EuanPyle/dynamo2relion). Particles were imported into an Alpha-phase development version of RELION 5.0 (4.1-alpha-1-commit- d2053c) and extracted as pseudo-subtomograms at bin4 (voxel size of 6.104 Å) in 64 voxel boxes. Pseudo-subtomograms are generated from the raw tilt-series and do not use denoised tomograms. A reference was reconstructed at the same box and voxel size using the Tomo Reconstruct Particle job-type. Particles were refined using Refine3D with the reference low-pass filtered to 30 Å, no mask applied, a particle diameter of 200 Å, and all Euler angles limited to local refinements of approximately 9 ° using the additional argument *--sigma_ang 3*. Poorly aligned particles were removed via 3D Classification without particle alignment, no mask applied, 6 classes, and a regularisation parameter (T value) of 0.2. A reference was reconstructed at bin1 in a box size of 196 voxels before the tilt series alignment for each tomogram was refined using Tomo Frame Alignment without fitting per- particle motion or deformations. Particles were re-extracted as pseudo-subtomograms at bin4 as before and refined as before using a mask over 1 inner coat subunit. The structure generated by RELION was used to pick more particles in CTF-corrected (non-denoised) tomograms with PyTOM template matching^44^ with dose-weighting and CTF- correction applied. The template used was filtered to 25 Å.

Coordinates from PyTOM (29496 particles from 475 tomograms) were imported into RELION 5.0. To remove junk particles, 3D classification was carried out with alignment using restricted Tilt and Psi Euler angles (*--sigma_rot 3 --sigma_psi 3*) but leaving in-plane rotation free, a mask over 1 inner coat unit and over part of neighbouring subunits, the map from the refined manually picked particle low-pass filtered to 25 Å as a reference, 4 classes, a T value of 0.1, and a particle diameter of 330 Å. Particles clearly resembling the COPII inner coat were kept, and refined under similar conditions to the preceding 3D classification. Particles were cleaned again using 3D classification without alignment with 6 classes and a T value of 0.2. The resulting particles.star file was merged with the manually picked particles generated earlier. Duplicate coordinates were deleted. Particles were exported to a Dynamo table using *relion2dynamo* (https://github.com/EuanPyle/relion2dynamo) and were cleaned by Neighbour Analysis, as previously described^47^. Coordinates were converted back to a .star file using *dynamo2relion* and reimported into RELION. Particles were refined as before, but at bin2 and with all Euler angles limited to local refinements using *--sigma_ang 3.* One more round of Tomo Frame Alignment, with per-particle motion, was carried out before Tomo CTF Refinement. A final refinement was carried out at bin2 from 12187 particles (from 352 tomograms), with limited Euler angles using *--sigma_ang 1.5.* The resolution according to Fourier Shell Correlation (FSC) (FSC = 0.143) was 14.4 Å (Extended Data Fig. 6A).

A difference map, as described in Figure 3A, between this structure and the inner coat from cargo-less GUVs was generated as follows: a model of the inner coat from cargo-less GUVs (PDB: 8BSH) was fitted into the inner coat map from microsomes. A volume representation of the fitted model was generated using *molmap* in UCSF Chimera^49^ at high resolution (2 Å) before low pass filtering to 14 Å in MATLAB. All maps were normalised to the same mean and standard deviation before the map from the fitted model was subtracted from our map from microsomes. Another difference map, as described in Figure 3B, was generated in the same way but using a 14 Å low pass filtered electron density map (EMDB-15949) corresponding to the fitted PDB model (PDB: 8BSH) instead of our map of the inner coat derived from microsomes.

##### Outer coat (vertex)

Outer coat vertices were manually picked in 30 tomograms. Particles were assigned Euler angles normal to their nearest membrane. Particles were extracted in 64 voxel boxes from RELION-reconstructed tomograms at a voxel size of 12.208 Å (bin8). Particles were averaged as before for the inner coat to form an initial average. Particles were then aligned and averaged in Dynamo as for the inner coat but with a translational shift of 4 voxels allowed and with a mask covering the vertex. The resulting map was used as a template to pick more particles in CTF-corrected tomograms with PyTOM template matching on all tomograms^44^, as for the inner coat. Particles were cleaned based on their proximity to the membrane of the vesicles. Particles were aligned in Dynamo again, and the resulting Dynamo table was converted to a .star file using *dynamo2relion*.

Vertex particles were imported into RELION 5.0 and extracted as pseudo-subtomograms in a box size of 64 voxels and at a voxel size of 6.104 Å (bin4). Particles were cleaned using 3D classification with refinement restricting the Tilt and Psi Euler angles (*--sigma_rot 4 -- sigma_psi 4*) but leaving in-plane rotation free. 3D classification used 3 classes, a T value of 0.25, and a particle diameter of 600 Å. Particles containing the vertex were then refined under the same conditions used in 3D classification. Particles were extracted at bin2 and further refined. Tomo frame alignment, Tomo Ctf refinement, and subsequent refinement at bin2 (voxel size of 3.052 Å) was iteratively repeated until resolution improvements stopped. The final map was reconstructed from 19368 particles and had a resolution of 11.4 Å (FSC = 0.143, Extended Data Fig. 6B).

##### Outer coat (rod)

Outer coat rods were manually picked in all tomograms. Particles were assigned Euler angles normal to their nearest membrane. Particles were extracted from Relion-reconstructed tomograms at bin8 (voxel size of 12.208 Å), averaged to form a reference, and aligned in Dynamo, as per the outer coat vertices. Rods of different length were selected and isolated using Neighbour Analysis. The resulting Dynamo table was converted to a .star file using *dynamo2relion*.

Rod particles were imported into RELION 5.0 and extracted as pseudo-subtomograms in a box size of 64 voxels at bin8. As for the outer coat vertices, particles were progressively unbinned from bin8 to bin2 (voxel size of 3.052 Å) and refined with restrictions to apply local Euler angle searches. The final map was reconstructed from 18852 particles and had a resolution of 11.8 Å (FSC = 0.143, Extended Data Fig. 6C).

To produce the maps in Figure 5D, particles were separated in different groups depending on the position of the neighbouring vertices as defined by masks on the neighbour plot. Classes contained between 1500 and 2500 particles each, were binned 8 times and were not filtetered beyond their Nyquist (at 24A).

##### Outer coat (vertex, 5-way rods)

We manually picked vertices formed by the convergence of 5 rods (as judged from visual inspection, see examples in Figure 4A). We extracted particles (n=461) from IsoNet corrected tomograms at bin8 (voxel size of 12.208 Å) in 64 voxel boxes, assigned initial angles normal to the membrane, and randomised the in-plane rotation, before averaging them to obtain a starting reference for alignments. We used dynamo to align particles by restraining the angles normal to the membrane withing a 20 degree cone, and allowing full searches for in plane rotation. After 50 iterations, alignments had converged. We imported the aligned coordinates in relion 4.1 and run a refinement at bin8, using angular restraints (--sigma_ang 3). The final map was reconstructed from 461 particles and had a resolution of 34 Å (FSC = 0.143).

#### GUVs-Sed5 Dataset

##### Inner coat

The surface of tubes in RELION-reconstructed tomograms was defined and segmented using the Pick Particle plug-in in Chimera as described previously^47,48^. The surface of the tube was oversampled, and coordinates were assigned Euler angles normal to the membrane. Particles were extracted in 32 voxel boxes from RELION-reconstructed tomograms at a voxel size of 10.8 Å. Particles were then aligned and averaged in Dynamo as before for the microsome inner coat dataset with several exceptions: in-plane rotation was restricted to 20 ° with azimuth flipping enabled; C1 symmetry was applied; particle translation was limited 15 voxels in all directions; alignment was carried out for 1 iteration.

Duplicates defined as particles within 4 voxels of another particle, were deleted with Dynamo’s separation in tomogram function during alignment. A previous inner coat structure (EMD-11199)^16^ was low-pass filtered and used as a reference. Particles were cleaned by Neighbour Analysis as before for the microsome inner coat dataset. The resulting Dynamo table was converted to a .star file using *dynamo2relion*.

Particles were imported into RELION 5.0 and extracted as pseudo-subtomograms at bin8. They were refined and progressively unbinned iteratively until bin1 before Tomo Frame Refinement and Tomo Ctf Refinement as previously described^16^. The final map was reconstructed from 178700 particles and had a resolution of 4.1 Å (FSC = 0.143, Extended Data Fig. 6D). The map was sharped using RELION’s LocalRes sharpening with a −50 B-factor.

##### Outer coat (vertex)

To pick outer coat vertices, we used the refined coordinates for the inner coat lattice and radially shifted them away from the membrane by 12 pixels. This was done to randomly oversample outer coat subunits at the expected radial distance from the tubular membrane. We then extracted these particles in a 64 voxel box size from RELION- reconstructed tomograms using Dynamo before aligning to a low pass filtered of a previous vertex structure (EMDB-11194)^15^. Alignment parameters were the same as was used for the inner coat alignment from the GUVs-Sed5 dataset except C2 symmetry was applied. The resulting Dynamo table was converted to a .star file using *dynamo2relion*.

Vertex particles were imported into RELION 5.0 and extracted as pseudo-subtomograms in a box size of 128 voxels at bin4. As for the outer coat vertices from microsomes, particles were progressively unbinned from bin8 to bin2 and refined with restrictions to apply local Euler angle searches. The final map was reconstructed from 13529 particles and had a resolution of 9.7 Å (FSC = 0.143, Extended Data Fig. 6E). The map was sharped using RELION’s LocalRes sharpening with a −175 B-factor.

##### Outer coat (rod)

To pick outer coat rods, we used the refined coordinates of the outer coat vertices and used Dynamo’s subboxing function to create 4 new coordinates where the rods are placed relative to each vertex. As before for the outer coat vertices, particles were aligned in Dynamo to a low pass filtered reference (EMDB-11193)^15^. Particles were cleaned by Neighbour Analysis and duplicates were deleted. The resulting Dynamo table was converted to a .star file using *dynamo2relion*.

Rod particles were imported into RELION 5.0 and extracted as pseudo-subtomograms in a box size of 128 voxels at bin4. As for the outer coat rods from microsomes, particles were progressively unbinned from bin8 to bin2 and refined with restrictions to apply local Euler angle searches. The final map was reconstructed from 39757 particles and had a resolution of 9.5 Å (FSC = 0.143, Extended Data Fig. 6F).

In all cases, relevant atomic model coordinates were rigid-body fitted into our maps using UCSF Chimera or ChimeraX. In all cases, fitting was unambiguous.

The number of particles for each dataset is summarised in Table 1.

**Table 1.**
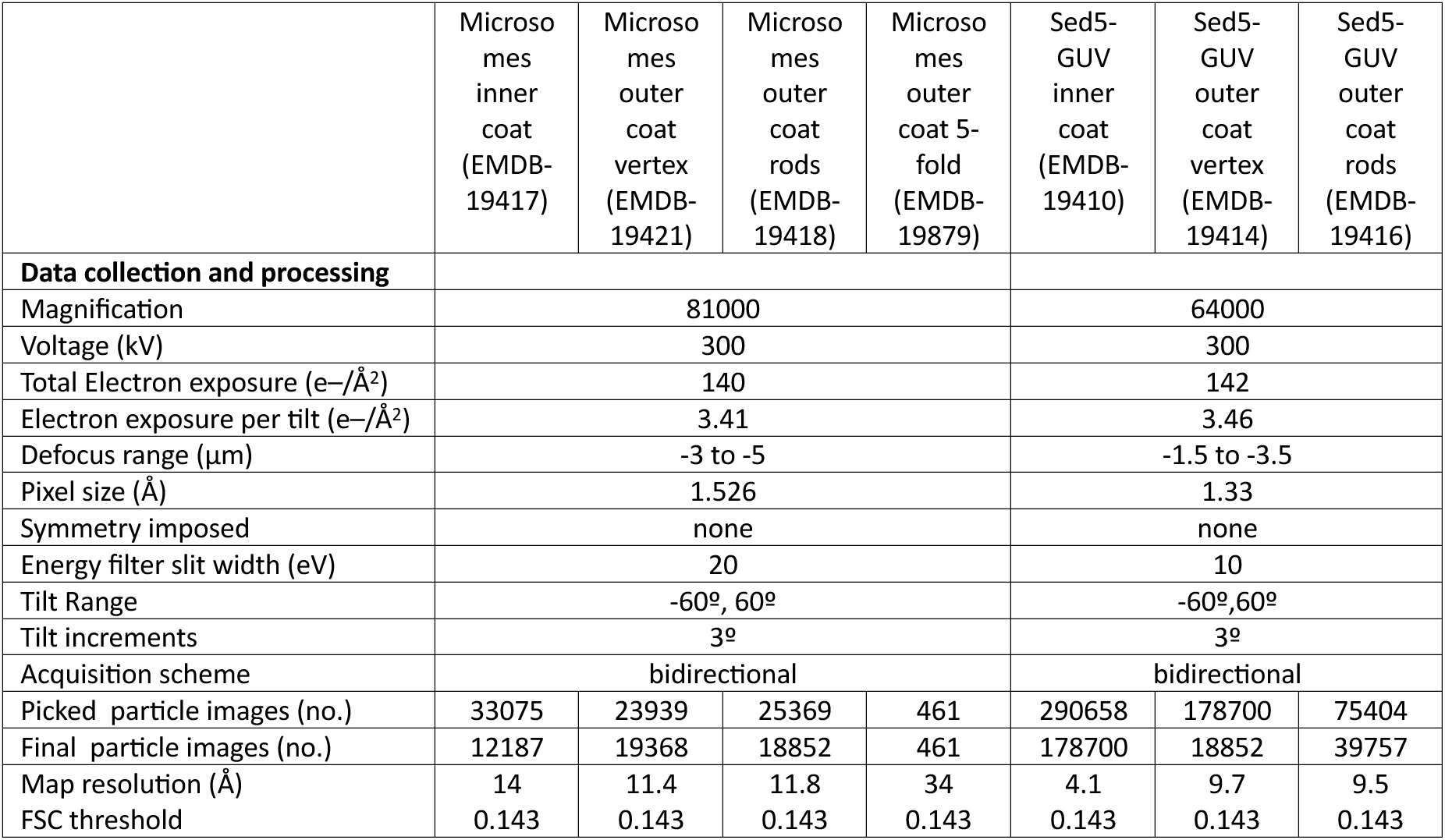
Cryo-EM data collecFon, refinement and validaFon staFsFcs.

|  | Microsomes inner coat (EMDB-19417) | Microsomes outer coat vertex (EMDB-19421) | Microsomes outer coat rods (EMDB-19418) | Microsomes outer coat 5-fold (EMDB-19879) | Sed5-GUV inner coat (EMDB-19410) | Sed5-GUV outer coat vertex (EMDB-19414) | Sed5-GUV outer coat rods (EMDB-19416) |
| --- | --- | --- | --- | --- | --- | --- | --- |
| **Data collection and processing** |  |  |  |  |  |  |  |
| Magnification | 81000 |  |  |  | 64000 |  |  |
| Voltage (kV) | 300 |  |  |  | 300 |  |  |
| Total Electron exposure (e-/Å<sup>2</sup>) | 140 |  |  |  | 142 |  |  |
| Electron exposure per tilt (e-/Å<sup>2</sup>) | 3.41 |  |  |  | 3.46 |  |  |
| Defocus range (µm) | -3 to -5 |  |  |  | -1.5 to -3.5 |  |  |
| Pixel size (Å) | 1.526 |  |  |  | 1.33 |  |  |
| Symmetry imposed | none |  |  |  | none |  |  |
| Energy filter slit width (eV) | 20 |  |  |  | 10 |  |  |
| Tilt Range | -60°, 60° |  |  |  | -60°, 60° |  |  |
| Tilt increments | 3° |  |  |  | 3° |  |  |
| Acquisition scheme | bidirectional |  |  |  | bidirectional |  |  |
| Picked particle images (no.) | 33075 | 23939 | 25369 | 461 | 290658 | 178700 | 75404 |
| Final particle images (no.) | 12187 | 19368 | 18852 | 461 | 178700 | 18852 | 39757 |
| Map resolution (Å) | 14 | 11.4 | 11.8 | 34 | 4.1 | 9.7 | 9.5 |
| FSC threshold | 0.143 | 0.143 | 0.143 | 0.143 | 0.143 | 0.143 | 0.143 |

### Mass spectrometry analysis

Total protein from the *S. cerevisiae* ER microsomes used in the reconstitution experiments (n=1) was digested using SP3 method^50^ with some adaptations. In brief, after reduction and alkylation of cysteines, total protein was precipitated onto magnetic beads (MagReSyn® Hydroxyl, Resyn Biosciences) by adding ethanol to a final concentration of 80% (v/v). Digestion was carried out by incubating the washed magnetic beads/total protein aggregated material with 1ug of trypsin (promega) dissolved in 25mM ammonium bicarbonate containing 0.1% RapiGest detergent (Waters). Sample was then acidified with trifluoroacetic acid to a final concentration of 0.5% (v/v) to stop digestion and induce RapiGest degradation. Magnetic beads and RapiGest in-soluble degradation products were pelleted by centrifugation at 11k xg for 15 min and the supernatant containing tryptic peptides was then taken for mass spectrometry analysis.

LC-MS/MS was performed on an Ultimate U3000 HPLC (ThermoFisher Scientific, San Jose, USA) hyphenated to an Orbitrap QExactive Classic mass spectrometer (ThermoFisher Scientific, San Jose, USA). Peptides were trapped on a C18 Acclaim PepMap 100 (5 µm, 300 µm x 5mm) trap column (ThermoFisher Scientific, San Jose, USA) and eluted onto a C18 Easy Spray Column, 2 µm, 75 µm x 500 mm (ThermoFisher Scientific, San Jose, USA) using 180 minute gradient of acetonitrile (5 to 40%). For data dependent acquisition, MS1 scans were acquired at a resolution of 70,000 (AGC target of 1e6 ions with a maximum injection time of 65ms) followed by ten MS2 scans acquired at a resolution of 17,500 (AGC target of 2e5 ions with a maximum injection time of 100ms) using a collision induced dissociation energy of 25. Dynamic exclusion of fragmented m/z values was set to 40s.

Raw data were imported and processed in Proteome Discoverer v3.1 (Thermo Fisher Scientific). The raw files were submitted to a database search using Proteome Discoverer with Sequest HT against the UniProt reference proteome for *S. cerevisiae.* Processing step consisted a double iterative search using INFERIS Rescoring algorithm on a first pass with methionine oxidation and cysteine carbamidomethylation set as variable and fixed modifications, repectively. For the second pass, all spectra with with a confidence filter worse than “high” were researched with Sequest HT including additional common protein variable modifications [Deamidated (N,Q); gln to pyro-Glu (Q); N-terminal acetylation and Methionine loss). The spectra identification was performed with the following parameters: MS accuracy, 10 p.p.m.; MS/MS accuracy of 0.02 Da; up to two trypsin missed cleavage sites were allowed. Percolator node was used for false discovery rate estimation and only rank 1 peptide identifications of high confidence (FDR<1%) were accepted.

## Data Availability

Data supporting the findings of this paper are available from the corresponding author upon reasonable request.

We have deposited the EM maps and models to the Electron Microscopy Data Bank with accession codes:

COPII Inner Coat on Microsome Vesicles: EMDB-19417

COPII Outer Coat (Rod) (Long) on Microsome Vesicles: EMDB-19421 COPII Outer Coat (Vertex) on Microsome Vesicles: EMDB-19418

COPII Outer coat (5-way vertex) on Microsome vesicles: EMDB-19879 COPII Inner Coat on Tubes from Sed5-GUVs: EMDB-19410

COPII Outer Coat (Rod) on Tubes from Sed5-GUVs: EMDB-19414 COPII Outer Coat (Vertex) on Tubes from Sed5-GUVs: EMDB-19416

## Supplementary Figures

**Supplementary Figure 1:**
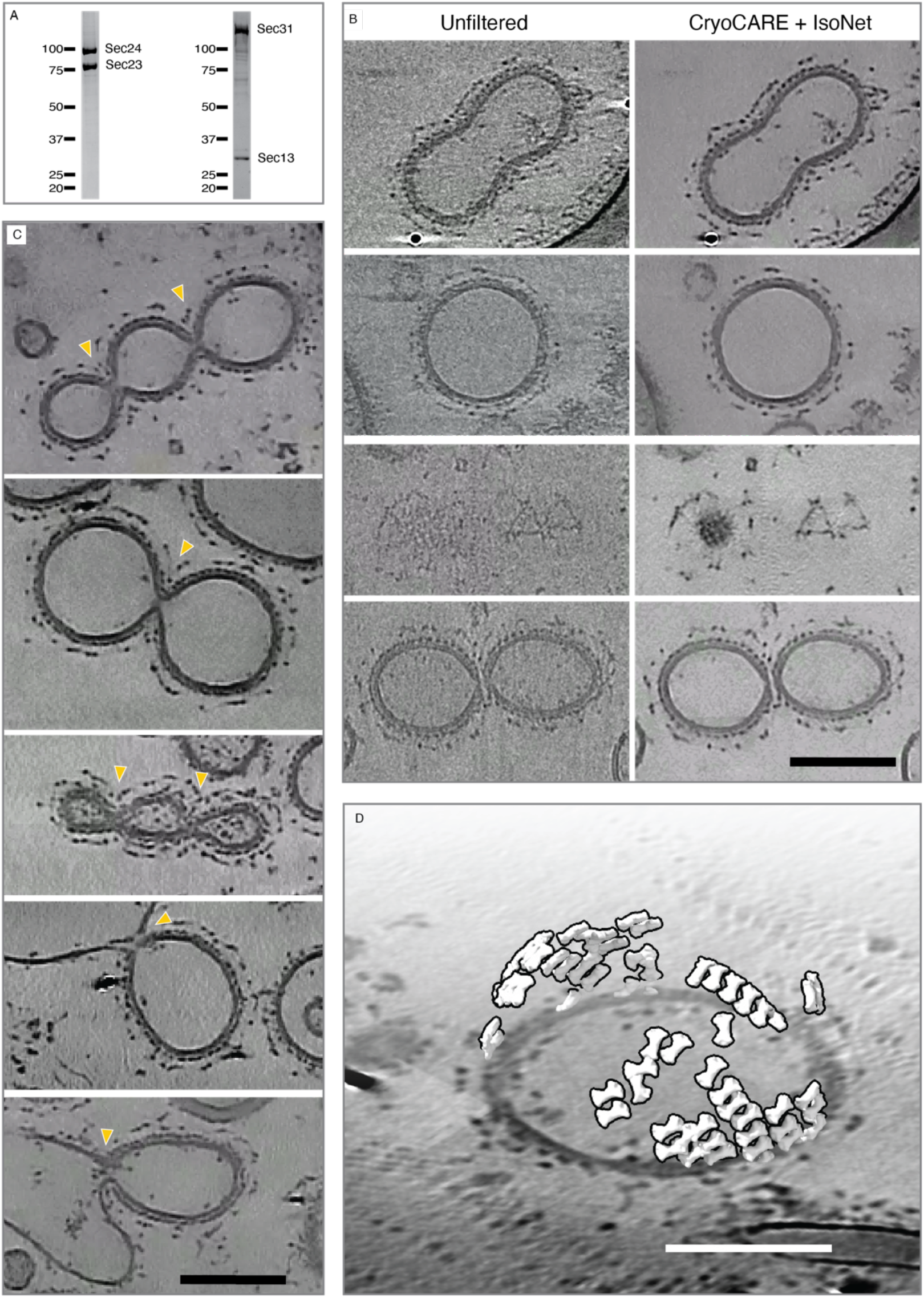
(A) Polyacrylamide gel of purified Sec23-Sec24 and Sec13-Sec31 complexes from insect cell expression. (B) XY slices through reconstructed tomograms comparing unfiltered 8X binned tomograms (left panels) to the corresponding IsoNet- treated ones (right panels) (C) XY slices through reconstructed tomograms that show rare events with constricted but not detached vesicle necks (yellow arrowheads). (D) Placed inner coat subunits overlayed to an orthoslice of a representative tomogram. Scale bars 100 nm.

**Supplementary Figure 2:**
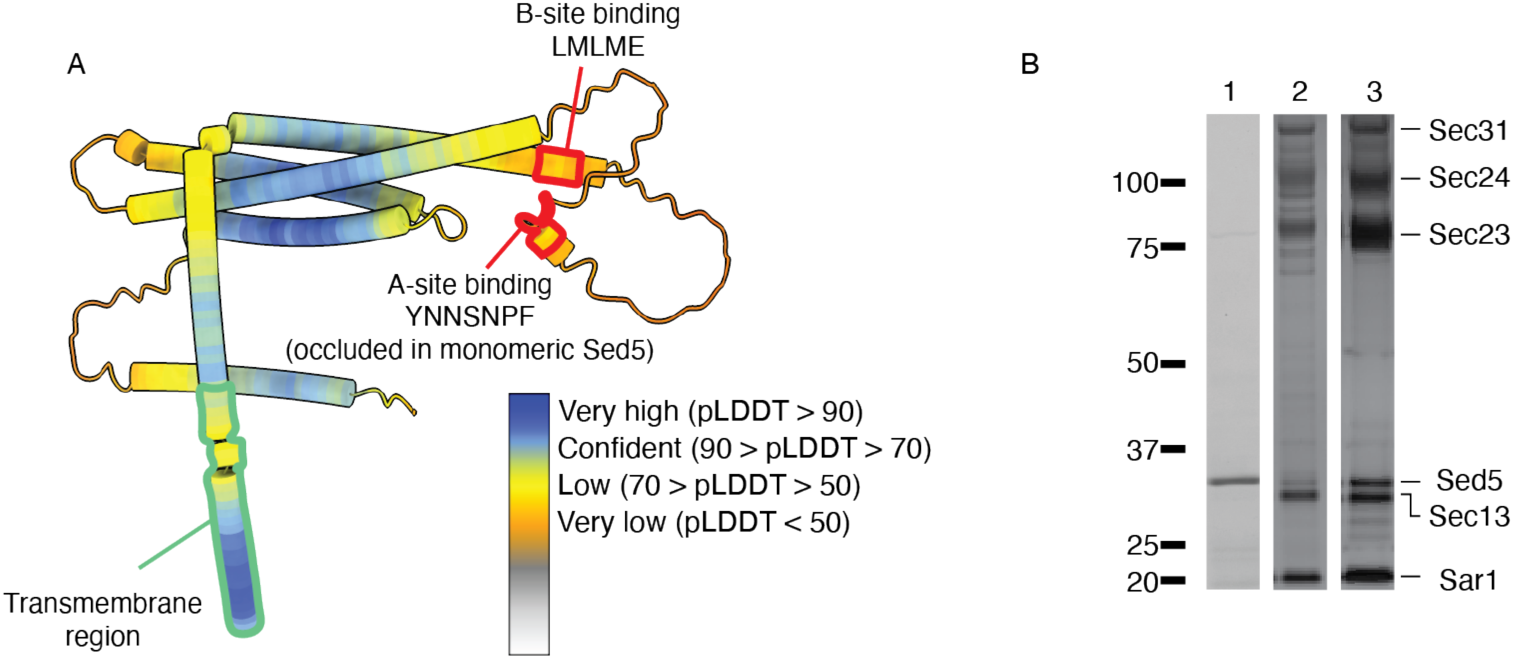
(A) Alpha-fold prediction of the full-length Sed5 structure (PDB ID: AF-Q01590-F1), coloured according to pLDDT value. The C-terminal transmembrane domain is highlighted in green, whilst the two Sec24-binding peptides are highlighted in red. These both fall within very low confidence regions, indicating disorder/flexibility. (B) Acrylamide gel showing purified Sed5 (lane 1), pelleted (lane 2) and floating (lane 3) fractions from a liposome flotation experiment.

**Supplementary Figure 3:**
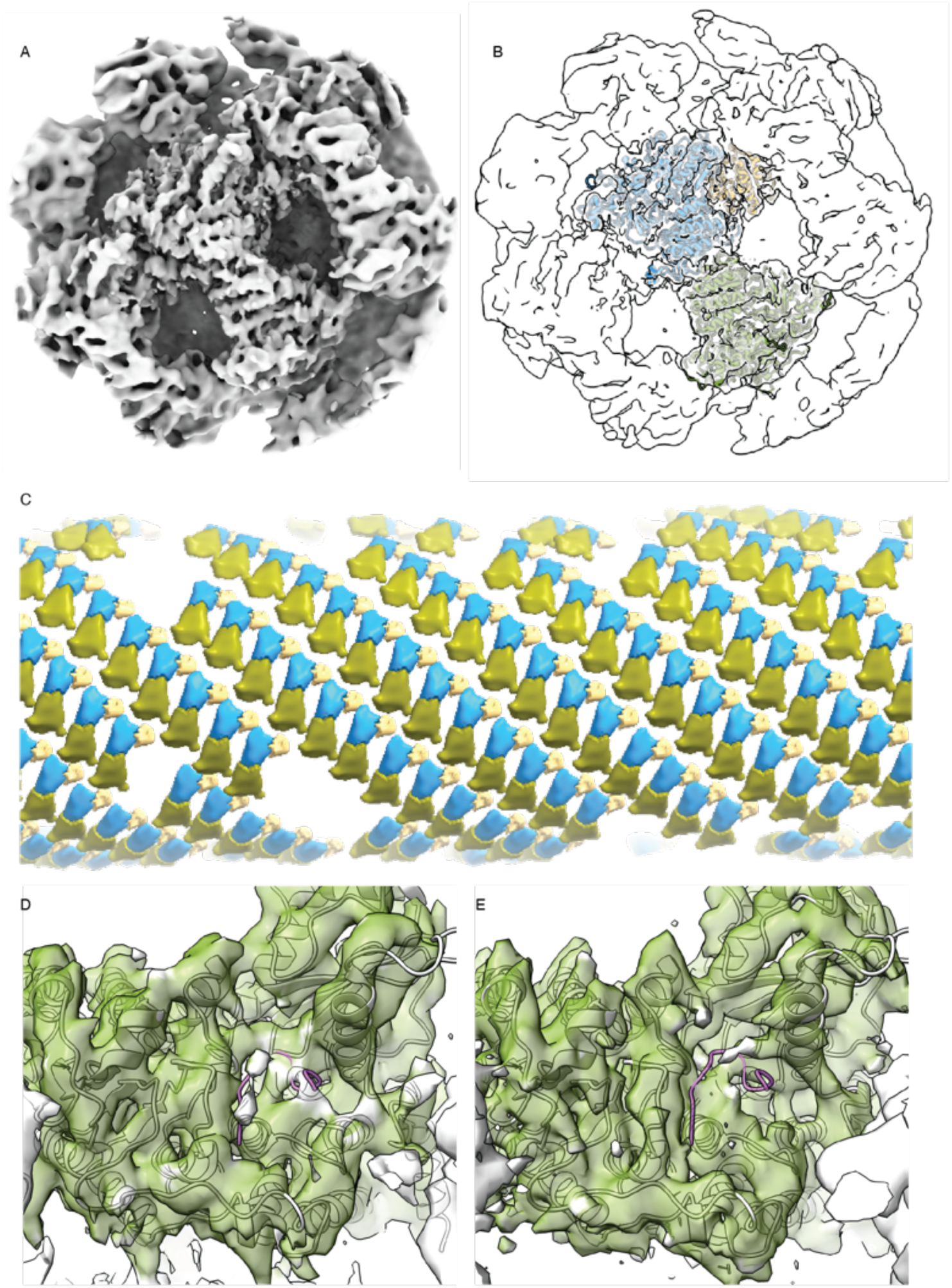
(A) Surface representation of the 4.1 Å STA map of the inner coat on Sed5-enriched GUV tubules (B) as in (A), with the Sec24-Sec24-Sar1 heterotrimer atomic model fitted. Sec23 in blue, Sec24 in green, Sar1 in yellow. (C) A 20 Å low-pass filtered map of the inner coat mapped back onto a representative section of a tomogram, showing the extensive lattice wrapping around a tubule. (D) and (E), As in Figure 3E and 3F, but focussing on the A-site of Sec24, and showing the Sed5 YNNSNPF peptide bound as crystallised (PDB ID: 8PD0, in purple). While extra density in (D) (Sed5-bound) with respect to (E) cannot be excluded, it is difficult to unambiguously detect above the noise.

**Supplementary Figure 4:**
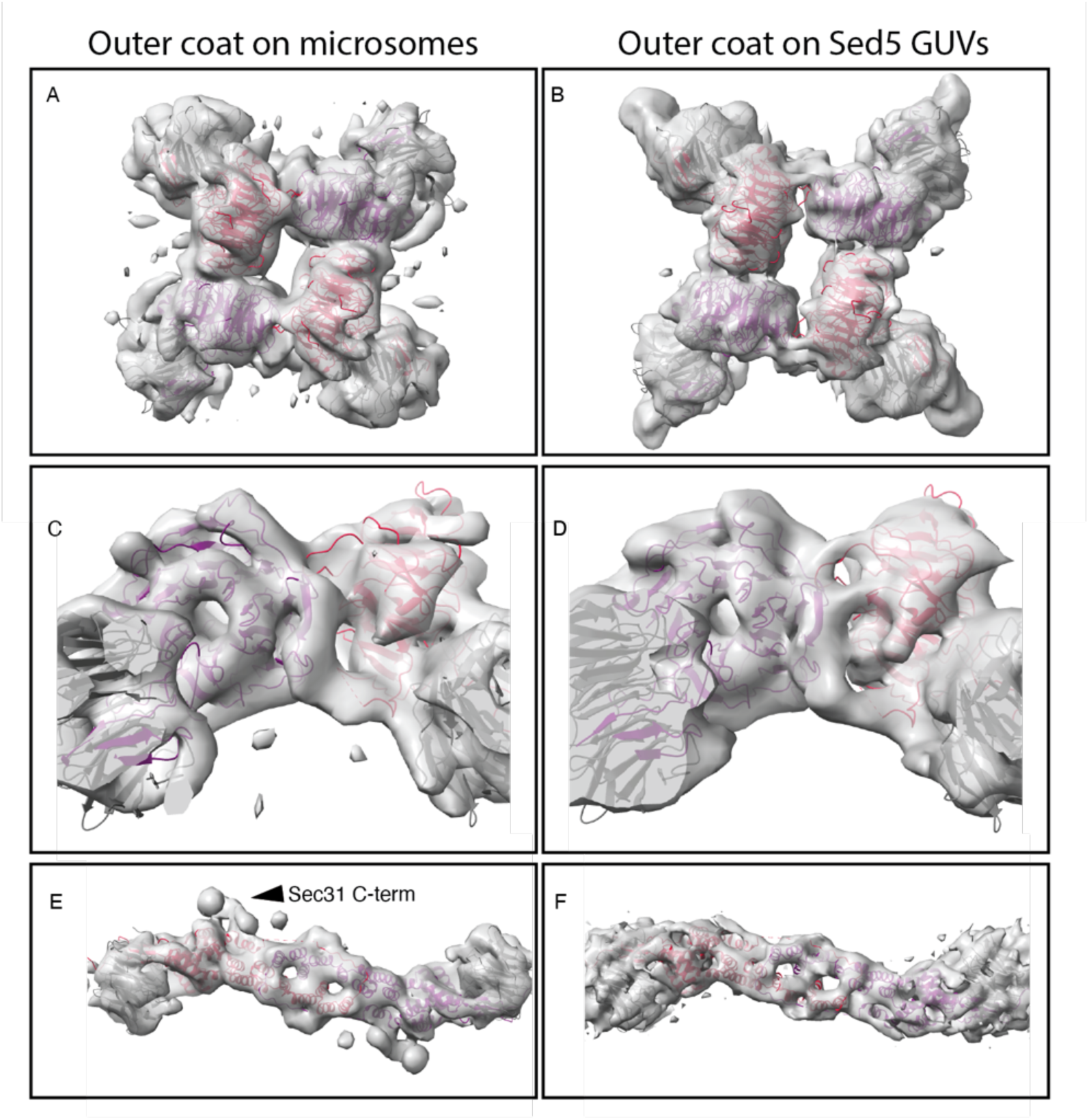
Comparison between the outer coat STA maps obtained from microsome and Sed5-GUV derived vesicles and tubes respectively. (A,B) overview of vertices, with four copies of the atomic model of the Sec13-Sec31 ‘vertex element’ fitted (PDB 2PM9). Sec31 in red and purple, Sec13 in grey. (C,D) close up of the vertices from a side view. (E,F), overview of rods, with the atomic model of the Sec13-Sec31 ‘edge’ element fitted (PDB 2PM6). Colour code as in (A,B).

**Supplementary Figure 5:**
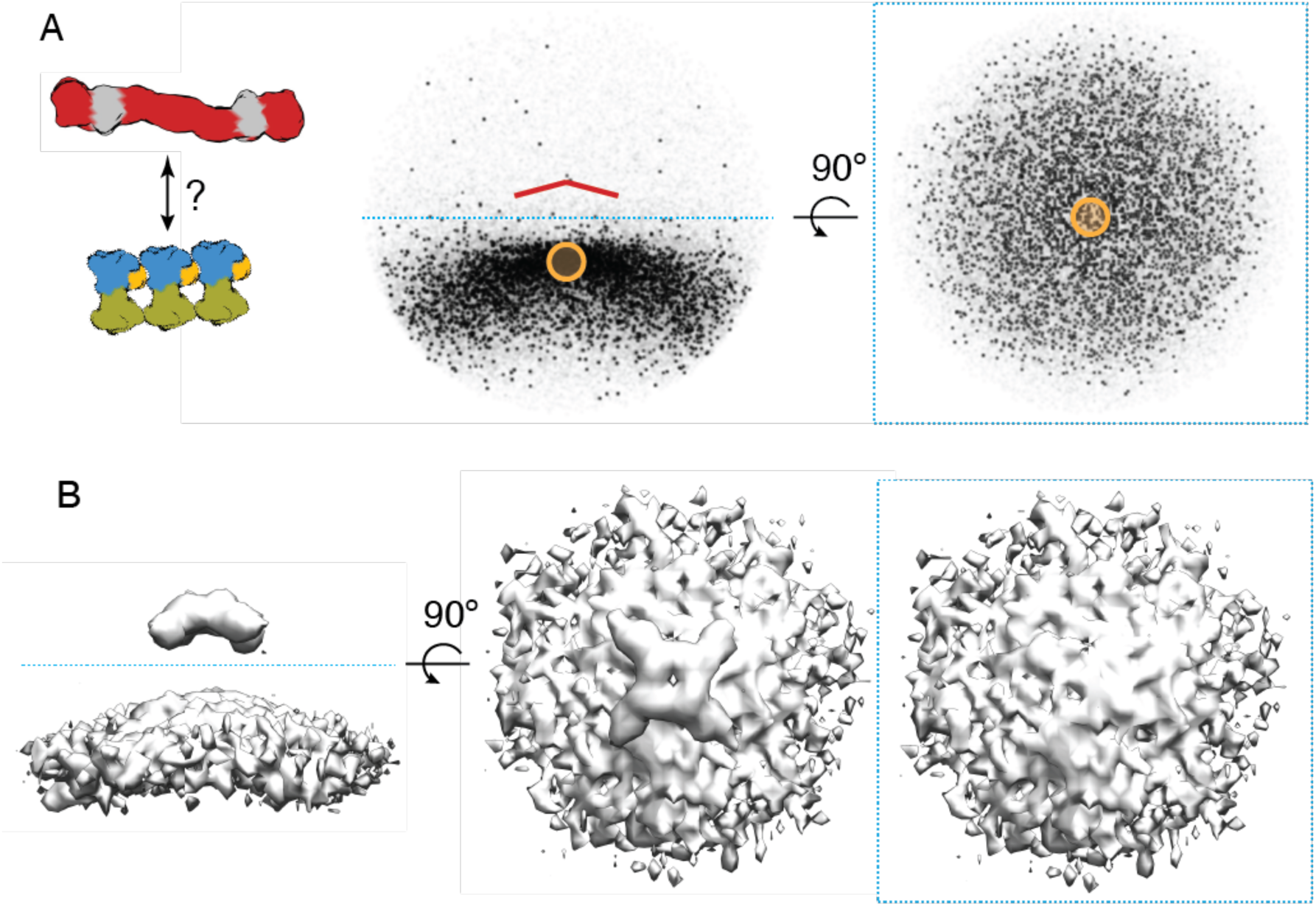
Relationship between inner and outer coat. A) Plot of positions of aligned inner coat subunits with respect to each aligned vertex, viewed from the side (middle panel), and from the top (right panel). Each black ‘dot’ represents an inner coat neighbour. While inner coat subunits are at expected positions ‘below’ vertices, the absence of a pattern suggest random translation between the two lattices B) Subtomogram average maps of vertices selected to have inner coat neighbours within the region shown in the yellow mask in (A), viewed from the side (left panel), from the top (middle panel), and from the top after removing the top half of the map (only showing density for the inner coat). The absence of any recognisable inner coat features indicated random rotation between the two lattices.

**Supplementary Figure 6:**
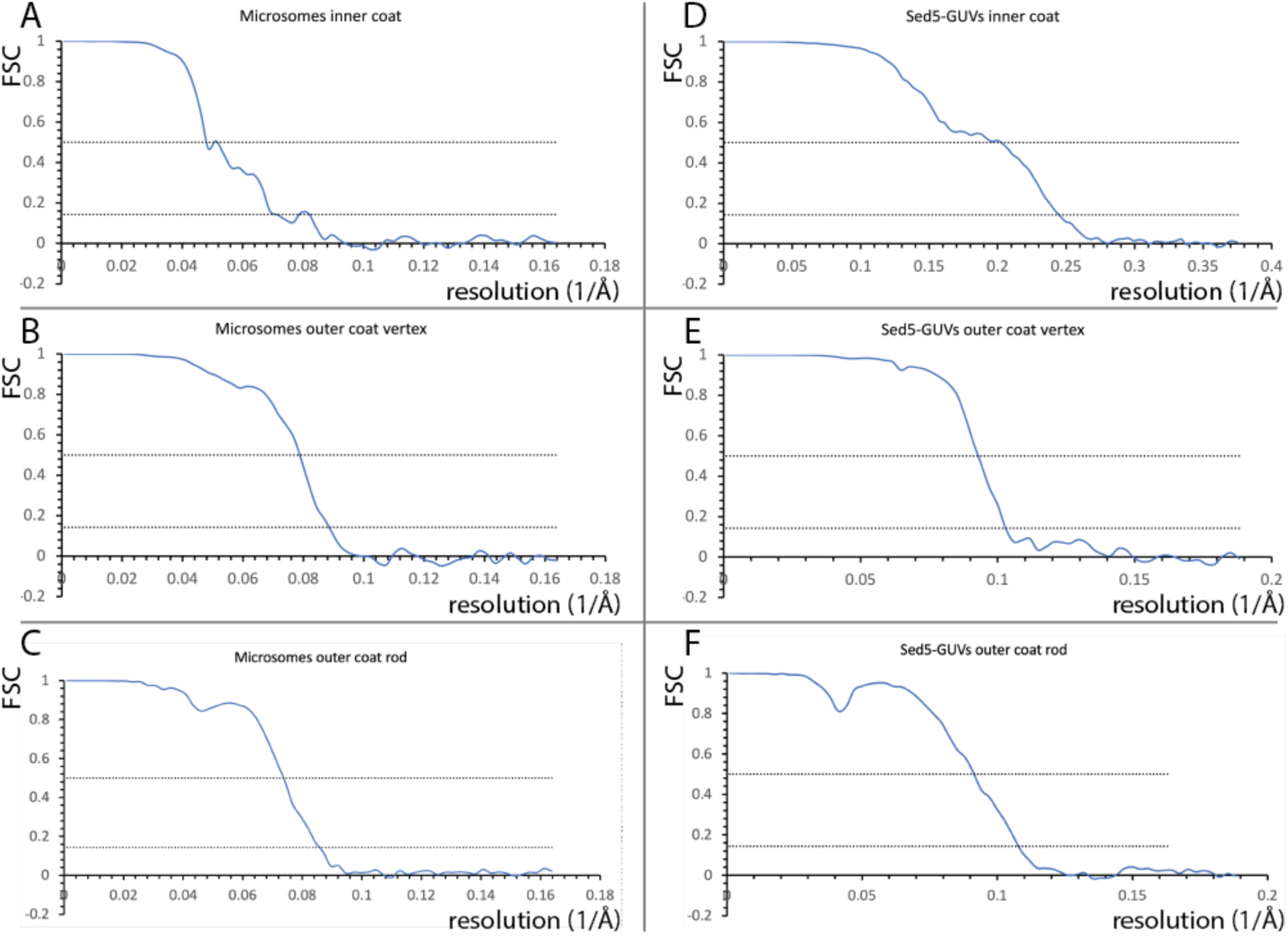
Fourier Shell Correlation plots for the maps discussed in this paper, as indicated by each chart title. 0.5 and 0.143 thresholds are shown as dotted lines for reference. All refinements were carried out with two independent halves, and the resolutions reported are measured at the 0.143 threshold.

## Notes

### Competing Interest Statement

The authors have declared no competing interest.

### Summary of Updates

We expanded the analysis of the outer coat layer, and generally updated the manuscript in response to peer review.

